# Deep Learning Prediction of DNA Damage Response Protein Signaling from H&E Histopathology and Reverse-Phase Protein Arrays in Pan-Gynecologic Cancers

**DOI:** 10.64898/2026.11.17.766760

**Authors:** Samantha Lane, Ying Zou

**Affiliations:** Bioinformatics, Johns Hopkins University; Johns Hopkins School of Medicine

## Abstract

Defects in homologous recombination and the DNA damage response (DDR) make gynecologic and breast tumors sensitive to platinum chemotherapy and PARP inhibitors, but identifying them requires costly genomic testing that reads permanent genomic scars rather than current pathway activity. Routine hematoxylin and eosin (H&E) slides are available for nearly every patient. This project asked whether H&E morphology carries information about functional DDR protein signaling across five cancer types in a single model.

We matched 1,673 diagnostic whole-slide images from The Cancer Genome Atlas (TCGA) to Level 4 reverse-phase protein array (RPPA) data for 1,527 patients with breast (BRCA, n = 858), endometrial (UCEC, n = 403), cervical (CESC, n = 146), ovarian (OV, n = 72), and uterine carcinosarcoma (UCS, n = 48) tumors. Slides were tiled at 0.5 µm/px and Macenko-normalized, yielding 20.4 million tiles. Tiles were embedded with two frozen pathology foundation models, UNI2-h and Virchow2. A gated attention-based multiple instance learning (ABMIL) network jointly regressed six RPPA pathway scores, including a DNA_damage composite of 11 DDR proteins. Performance was assessed by out-of-fold Spearman correlation under 5-fold, cancer-type-stratified, site-grouped cross-validation and compared against a baseline that predicts each patient’s cancer-type mean.

Predicted DNA_damage correlated with measured values (ρ = 0.43 for both encoders) and exceeded the cancer-type baseline (ρ = 0.31; Δρ = +0.13). The signal held within cancer types (within-type ρ = 0.29 for UNI2-h, 0.30 for Virchow2). With the original UNI encoder, it survived site-held-out folds with residualized targets (within-type ρ = 0.26), and it vanished when targets were shuffled within cancer type (within-type ρ ≤ 0.00 for all three encoders). All six pathway scores were predicted above baseline; Cell_cycle was strongest (ρ ≈ 0.59). In interpretability analyses, the top 10 attention tiles per slide contained more tumor (+1.8 SD), larger tumor nuclei (+1.0 to +1.3 SD), and fewer lymphocytes (−0.6 SD) than the bottom 10 tiles; 13 of 14 morphometric features differed (q < 0.05). Attention agreed with Integrated Gradients (median ρ ≈ 0.70).

Routine H&E contains a moderate, reproducible signal of DDR protein abundance beyond cancer type, concentrated in tumor-dense regions with enlarged nuclei. Separating this signal from proliferation and predicting phosphorylated CHK1 and CHK2 directly are the next steps toward a functional, slide-based DDR biomarker.

## Background and Significance

Existing H&E deep learning models predict static genomic alterations in one organ at a time; this project tested whether a single model can predict DDR protein signaling across five gynecologic and breast cancer types. That distinction matters because a mutation or genomic scar does not guarantee that the pathway is failing today, and treatment response depends on current pathway function.

Cells sense DNA double-strand breaks through the ATM kinase, which phosphorylates CHK2 at threonine 68, and sense replication stress through ATR, which phosphorylates CHK1 at serine 345 (Jackson & Bartek, 2009). These cascades trigger checkpoint arrest and homologous recombination (HR) repair through BRCA1, BRCA2, and RAD51. Tumors with HR deficiency (HRD) depend on alternative repair and are selectively killed by platinum agents and poly(ADP-ribose) polymerase (PARP) inhibitors (Lord & Ashworth, 2016).

Clinical trials show the size of this effect. In the TNT trial, carboplatin produced a 68% objective response rate in triple-negative breast cancer patients with germline BRCA1/2 mutations, compared with 33% for docetaxel in the same subgroup (Tutt et al., 2018). In OlympiA, adjuvant olaparib reduced invasive disease recurrence or death by 42% in germline BRCA1/2 carriers (Tutt et al., 2021).

Current HRD assays score loss of heterozygosity, telomeric allelic imbalance, and large-scale state transitions. These are permanent footprints of past repair failure. Tumors can restore HR through BRCA1/2 reversion mutations or promoter demethylation and still test HRD-positive, which helps explain why genomic HRD imperfectly predicts response (Lord & Ashworth, 2016). Functional assays such as RAD51 foci scoring track response better but require specialized immunofluorescence on fresh material (Castroviejo-Bermejo et al., 2018). Sequencing-based tests also add weeks of turnaround and cost, and fail on inadequate tissue (Bergstrom et al., 2024).

H&E slides are prepared for nearly every tumor biopsy at low cost (Koivukoski et al., 2023). Chromosomal instability alters nuclear size, shape, and chromatin texture in ways visible on these slides (Fernandes, 2024). DeepHRD, a weakly supervised model, predicts genomic HRD and platinum response from H&E in breast and ovarian cancer (Bergstrom et al., 2024).

Vision transformer models combined with nuclear morphometry infer TP53 and BRCA1 status in ovarian cancer (Fernandes, 2024). Both predict genomic labels, and both were developed within single organs.

Reverse-phase protein arrays quantify total and phosphorylated proteins across hundreds of samples. The Cancer Proteome Atlas (TCPA) provides normalized RPPA data for TCGA tumors (Li et al., 2013; Chen et al., 2019). Pan-cancer analysis of these data defined pathway scores, including a DNA damage response score built from 11 DDR proteins (Akbani et al., 2014). These scores give a continuous, protein-level training target that sits closer to pathway function than mutation status.

No prior pipeline has predicted continuous RPPA pathway activity from H&E, and none has done so across a pooled pan-gynecologic cohort. Pathology foundation models now make this feasible without manual annotation. UNI was pretrained on over 100 million images from more than 100,000 H&E slides (Chen et al., 2024), and its successor UNI2-h on over 200 million tiles from more than 350,000 H&E and IHC slides (MahmoodLab, n.d.). Virchow2 was pretrained on millions of slides at mixed magnifications (Zimmermann et al., 2024). Combined with attention-based multiple instance learning (Ilse et al., 2018), these models can map whole slides to patient-level continuous targets and indicate which regions drove each prediction.

The central hypothesis was that nuclear atypia, chromatin changes, and tumor microenvironment architecture reflect DDR signaling that deep networks can detect on routine H&E. The project had three aims: (1) assemble a matched H&E and RPPA cohort across five TCGA cancer types; (2) predict continuous RPPA pathway scores, with DNA_damage as the primary endpoint; and (3) characterize the morphology underlying the predictions. The long-term goal is a slide-based functional biomarker that could help prioritize patients for confirmatory DDR testing; clinical use would require validation well beyond this study.

## Materials and Methods

The pipeline matched TCGA slides to RPPA by patient, tiled and normalized each slide, embedded tiles with frozen foundation models, and trained an attention-based regression model under site-grouped cross-validation. Jobs ran as Slurm batch jobs using Python and PyTorch; feature extraction, model training, HoVer-Net, and Integrated Gradients ran on a Tesla T4 GPU, and the remaining steps on CPU nodes.

Diagnostic formalin-fixed, paraffin-embedded (FFPE) H&E whole-slide images (WSIs) were downloaded from the NCI Genomic Data Commons (GDC; Grossman et al., 2016) for five TCGA projects: BRCA, UCEC, CESC, OV, and UCS. Level 4 replicate-based normalized (RBN) RPPA data were obtained from the TCGA pan-cancer RPPA release (file

TCGA-RPPA-pancan-clean; Li et al., 2013; Chen et al., 2019). Slides and RPPA profiles were joined on patient barcode and sample type, so both came from the same specimen type. Patients without a usable diagnostic slide or RPPA profile were excluded. All slides from one patient formed a single bag for modeling.

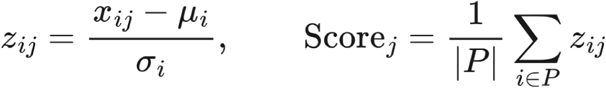

Six pathway scores were computed following the TCGA pan-cancer definitions (Akbani et al., 2014): DNA_damage, Cell_cycle, TSC_mTOR, PI3K_AKT, RAS_MAPK, and RTK. Each protein was z-scored using its mean and SD across all patients in the TCGA pan-cancer RPPA file (one profile per patient), so 0 is the pan-cancer average. A pathway score was set to missing if fewer than half of its member proteins were measured.

Here, x is the Level 4 log2 level of protein i in patient j, and P is the pathway’s member set. The DNA_damage score averages 11 proteins: nine total proteins (ATM, MRE11, RAD50, BRCA2, RAD51, KU80, 53BP1, P53, XRCC1) and two phosphoproteins (CHK1_pS345, CHK2_pT68). Because nine of the 11 components are total proteins, the score mainly reflects the abundance of repair machinery rather than kinase activity. For training, targets were re-standardized using training-fold statistics, and missing values were masked from the loss.

Slides were resampled to 0.5 µm per pixel (approximately 20x magnification). Tissue was segmented with an Otsu threshold on the saturation channel, and non-overlapping 256×256-pixel tiles with at least 50% tissue were extracted. Stain variation was reduced with Macenko normalization, mapping each slide’s stain vectors to a shared reference (Macenko et al., 2009). Quality control checked for consistent resolution and tiling, complete and non-duplicated bags, and outliers in stain, tissue area, and tile count.

Each tile was embedded once with two frozen encoders: UNI2-h, which outputs 1,536-dimensional features (MahmoodLab, n.d.), and Virchow2, which outputs 2,560-dimensional features (Zimmermann et al., 2024). An earlier run used the original UNI (ViT-L/16, 1,024 dimensions) and is reported as a robustness check. Features were cached to disk so that model training did not repeat the embedding step.

A gated attention ABMIL model (Ilse et al., 2018) mapped each patient’s bag of tile embeddings to six continuous outputs. Tile embeddings (h_k_) were projected to a 512-dimensional hidden layer. Gated attention weights were computed as:

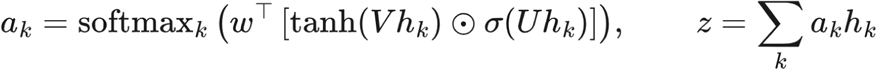

The attention layer had 256 units. The pooled slide representation z fed a two-layer regression head (512 → 256 → 6, ReLU) with six outputs, trained jointly. Dropout was 0.25. The model was trained with mean squared error loss, the AdamW optimizer (learning rate 1×10^-4^, weight decay 1×10^-4^) with cosine learning-rate decay, gradient accumulation over 8 bags, and bfloat16 mixed precision. Bags larger than 8,000 tiles were randomly subsampled at each training step; all tiles were used at evaluation. Training ran for up to 50 epochs with early stopping (patience 10), keeping the checkpoint with the lowest validation loss. Full settings are listed in Appendix A.

Patients were split into five folds, stratified by cancer type and grouped by TCGA tissue source site so that no site appeared in both training and test data. For each test fold, the next fold served as the validation set for early stopping and the remaining three folds were used for training. Each patient received exactly one out-of-fold prediction. The primary metric was the pooled out-of-fold Spearman correlation (ρ) between predicted and measured scores, with 95% confidence intervals from bootstrap resampling of patients. P-values were adjusted with the Benjamini-Hochberg procedure (Benjamini & Hochberg, 1995).

Pooled correlations can be inflated by differences between cancer types alone. Each model was therefore compared with a baseline that predicts each patient’s cancer-type mean from the training folds. Gain over this baseline is reported as Δρ. Within-type ρ was computed by centering measured and predicted scores on their cancer-type means and calculating one Spearman correlation across all patients. Aim 2 was considered successful for an endpoint if its 95% CI excluded zero, it beat the cancer-type baseline, and it remained positive within cancer types.

Two additional runs tested whether the model was learning cancer type or site artifacts. In the residual-target run, each target was residualized against cancer type and the model was trained with tissue-source-site-held-out folds. In the null (control) run, target values were shuffled within each cancer type, which preserves between-type differences but removes patient-level signals. The residual-target run used the original UNI encoder. Null runs were performed with UNI (residualized targets, site-held-out folds) and with UNI2-h and Virchow2 (standard targets and folds).

For each held-out slide, the 10 tiles with the highest attention and the 10 with the lowest were selected. Nuclei in these tiles were segmented and classified with HoVer-Net (Graham et al., 2019). Fourteen features were computed per tile, including tumor fraction, lymphocyte fraction, tumor nuclear area, and chromatin homogeneity. For each patient, the difference between median feature values in high- and low-attention tiles was averaged across slides.

Differences were tested with a two-sided Wilcoxon signed-rank test across patients with Benjamini-Hochberg (BH) correction, and expressed as the mean difference divided by the feature’s SD across selected tiles. “Lymphocytes” refers to HoVer-Net’s PanNuke inflammatory class, and mitoses were not quantified. Integrated Gradients (Sundararajan et al., 2017) were computed for the DNA_damage output with respect to each tile’s embedding. Agreement between attention and absolute attribution was measured by the per-slide Spearman correlation and summarized by its median. Tile-quality metrics, such as blur, tissue fraction, pen marks, etc., were computed and tested in the same way.

## Results

H&E predicted the DNA_damage score above a cancer-type baseline with both encoders (ρ = 0.43 vs 0.31), and the signal held within cancer types, across held-out sites, and against a permutation null. High-attention regions were tumor-dense, with enlarged nuclei and few lymphocytes.

### Aim 1 A matched pan-gynecologic cohort

The final cohort contained 1,527 patients with 1,673 diagnostic slides, fewer than the approximately 1,800 patients projected in the proposal. The proposal’s estimate counted all GDC slides, including frozen-section slides. Restricting to diagnostic FFPE slides with an RPPA profile from the same specimen type left 1,527 patients, and about 30 slides were not processed because of a missing file or read error. Tiling yielded ∼20.4 million tissue tiles. Breast cancer made up more than half of the cohort, and OV and UCS together contributed under 8% (Table 1).

**Table 1.** Matched cohort by cancer type.

| **TCGA project** | **Tissue** | **Patients** | **Share of cohort** |
|---|---|---|---|
| BRCA | Breast | 858 | 56.2% |
| UCEC | Endometrial | 403 | 26.4% |
| CESC | Cervical | 146 | 9.6% |
| OV | Ovarian | 72 | 4.7% |
| UCS | Uterine carcinosarcoma | 48 | 3.1% |
| Total |  | 1,527 |  |

### Aim 2 H&E predicts RPPA pathway scores beyond cancer type

For the primary endpoint, DNA_damage, the UNI2-h model reached ρ = 0.43 (95% CI 0.39-0.47) and the Virchow2 model ρ = 0.43 (0.39-0.48). Both exceeded the cancer-type baseline of ρ = 0.31 by Δρ = +0.13. Within cancer types, ρ was 0.29 for UNI2-h and 0.30 for Virchow2.

With UNI2-h, DNA_damage predictions were significant within all five cancer types (ρ = 0.25-0.48). With Virchow2, CESC was borderline (ρ = 0.16, 95% CI −0.00 to 0.32, q = 0.066), so only UNI2-h strictly met the within-type success criterion. Per-type values appear in Table 2.

**Table 2.** DNA_damage Spearman ρ within each cancer type.

| Cancer type | n | UNI2-h ρ (95% CI) | Virchow2 ρ (95% CI) |
|---|---|---|---|
| BRCA | 858 | 0.27 (0.21 to 0.33) | 0.31 (0.25 to 0.36) |
| UCEC | 403 | 0.31 (0.22 to 0.40) | 0.31 (0.22 to 0.40) |
| CESC | 146 | 0.25 (0.09 to 0.39) | 0.16 (−0.00 to 0.32) |
| OV | 72 | 0.48 (0.27 to 0.64) | 0.45 (0.25 to 0.61) |
| UCS | 48 | 0.38 (0.12 to 0.58) | 0.39 (0.16 to 0.58) |

All six pathway scores were predicted above baseline, with within-type ρ ranging from 0.28 to 0.48 across pathways. Cell_cycle was the strongest target (pooled ρ = 0.59 for UNI2-h and 0.60 for Virchow2, against a cancer-type baseline of 0.33; within-type ρ = 0.46 and 0.48). The two encoders performed almost identically on every target, with overlapping confidence intervals throughout. Pooled R^2^ ranged from 0.12 to 0.32, and MSE from 0.12 to 0.25 (Table 3). R^2^ exceeded the cancer-type baseline for every target, with 95% CIs for ΔR^2^ excluding zero. However, the gain was modest for DNA_damage (ΔR^2^ ≈ +0.05), whose baseline R^2^ (0.17) was the highest of any target. Cell_cycle showed the largest gain (ΔR^2^ ≈ +0.18).

**Table 3.**
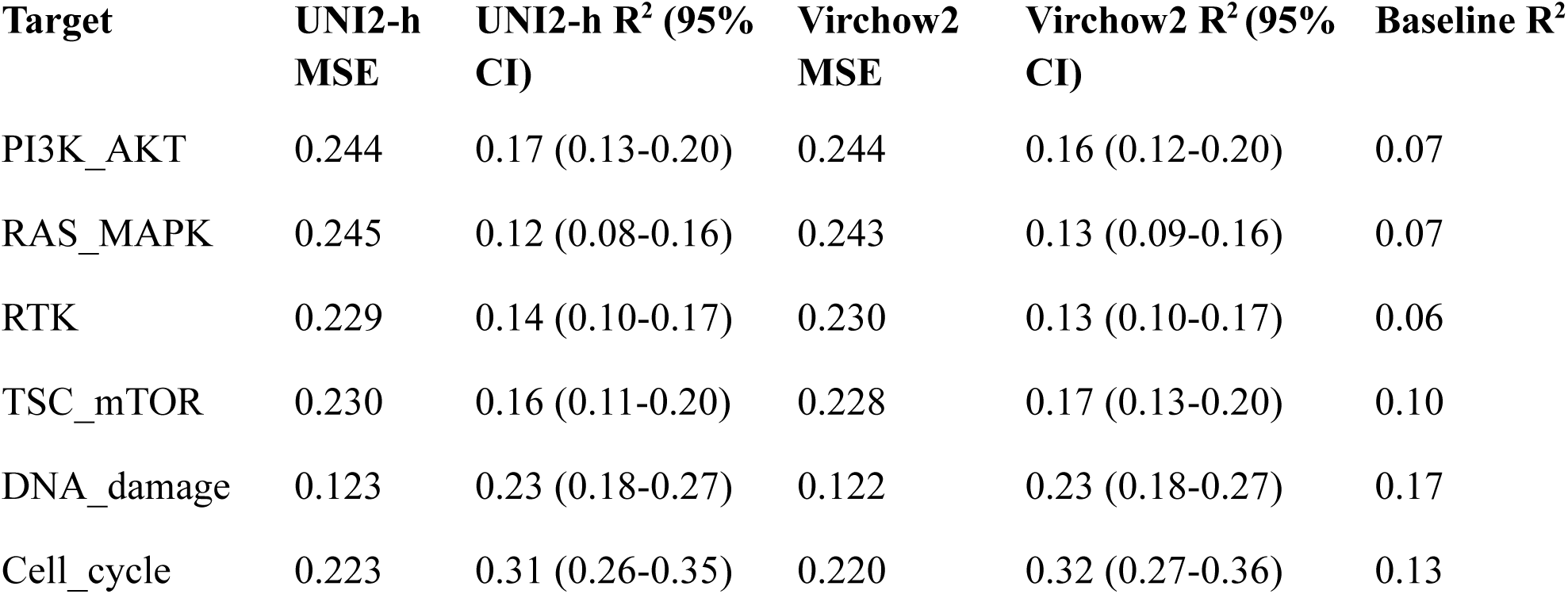
Pooled regression performance per target (n = 1,527; out-of-fold predictions, 5-fold site-grouped CV). MSE = RMSE²; R² shown with 95% bootstrap CI; baseline predicts each cancer type’s mean.

| **Target** | **UNI2-h MSE** | **UNI2-h R<sup>2</sup> (95% CI)** | **Virchow2 MSE** | **Virchow2 R<sup>2</sup> (95% CI)** | **Baseline R<sup>2</sup>** |
|---|---|---|---|---|---|
| PI3K_AKT | 0.244 | 0.17 (0.13-0.20) | 0.244 | 0.16 (0.12-0.20) | 0.07 |
| RAS_MAPK | 0.245 | 0.12 (0.08-0.16) | 0.243 | 0.13 (0.09-0.16) | 0.07 |
| RTK | 0.229 | 0.14 (0.10-0.17) | 0.230 | 0.13 (0.10-0.17) | 0.06 |
| TSC_mTOR | 0.230 | 0.16 (0.11-0.20) | 0.228 | 0.17 (0.13-0.20) | 0.10 |
| DNA_damage | 0.123 | 0.23 (0.18-0.27) | 0.122 | 0.23 (0.18-0.27) | 0.17 |
| Cell_cycle | 0.223 | 0.31 (0.26-0.35) | 0.220 | 0.32 (0.27-0.36) | 0.13 |

### Aim 2 Robustness to site, encoder, and permutation

The DNA_damage signal changed little when whole tissue source sites were held out and targets were residualized against cancer type, and it disappeared when targets were shuffled within cancer type (Table 4).

**Table 4.** DNA_damage robustness checks (within-type Spearman ρ).

| **Run** | **Encoder** | **Within-type ρ (95% CI)** | **Δρ vs baseline** |
|---|---|---|---|
| Standard CV (site-grouped) | UNI2-h | 0.29 (0.25 to 0.34) | +0.13 |
| Standard CV (site-grouped) | Virchow2 | 0.30 (0.26 to 0.35) | +0.13 |
| Site-held-out, residualized targets | UNI | 0.26 (0.21 to 0.30) | +0.12 |
| Targets shuffled within type | UNI (residualized) | 0.00 (−0.05 to 0.05) | +0.01 |
| Targets shuffled within type | UNI2-h | −0.02 (−0.08 to 0.02) | −0.04 |
| Targets shuffled within type | Virchow2 | −0.03 (−0.08 to 0.02) | −0.04 |

The shuffled run still showed a raw pooled ρ of 0.32. This is expected: shuffling within type preserves between-type differences, so the model can only reproduce the type-mean baseline (ρ = 0.31). In the residualized run, DNA_damage remained significant within BRCA, UCEC, and CESC; OV was borderline (q = 0.058) and UCS was not significant (ρ = 0.15, n = 48), likely reflecting the small size of those cohorts.

An earlier run with the original UNI encoder gave a slightly higher pooled DNA_damage ρ of 0.46 (0.42-0.50), Δρ = +0.16, with per-type ρ of 0.29-0.37. That run used patient-level rather than site-grouped folds: 76 of 110 tissue source sites contributed patients to more than one fold. Because TCGA sites differ in staining and scanning, site leakage may have inflated this estimate. When the same encoder was evaluated with site-held-out folds, within-type ρ fell to 0.26, in line with the UNI2-h and Virchow2 results.

### Aim 3 What the model attends to

Compared with the 10 lowest-attention tiles on each slide, the 10 highest-attention tiles contained substantially more tumor, larger tumor nuclei, and fewer lymphocytes. Thirteen of 14 morphometric features differed significantly (q < 0.05) under both encoders (Table 5).

**Table 5.** Selected morphometric differences, top-10 vs bottom-10 attention tiles (standard deviation units).

| **Feature** | **UNI2-h (SD)** | **q** | **Virchow2 (SD)** | **q** |
|---|---|---|---|---|
| Tumor fraction | +1.80 | 3.8e−248 | +1.86 | 2.1e−249 |
| Tumor nuclear area | +1.26 | 5.6e−190 | +1.03 | 6.3e−142 |
| Chromatin homogeneity | −0.42 | 1.2e−60 | −0.61 | 4.6e−66 |
| Lymphocyte fraction | −0.62 | 1.3e−155 | −0.61 | 8.1e−138 |

13 of 14 features differed under each encoder; the exception was chromatin hematoxylin SD for UNI2-h and chromatin entropy for Virchow2 (Appendix C).

Attention agreed with absolute Integrated Gradients attribution for DNA_damage (median per-slide ρ ≈ 0.70). Figure 2 shows both maps for an example held-out ovarian slide. Tile-quality metrics also differed between high- and low-attention tiles, so part of the morphological contrast may reflect image quality rather than biology.

**Figure 1.**
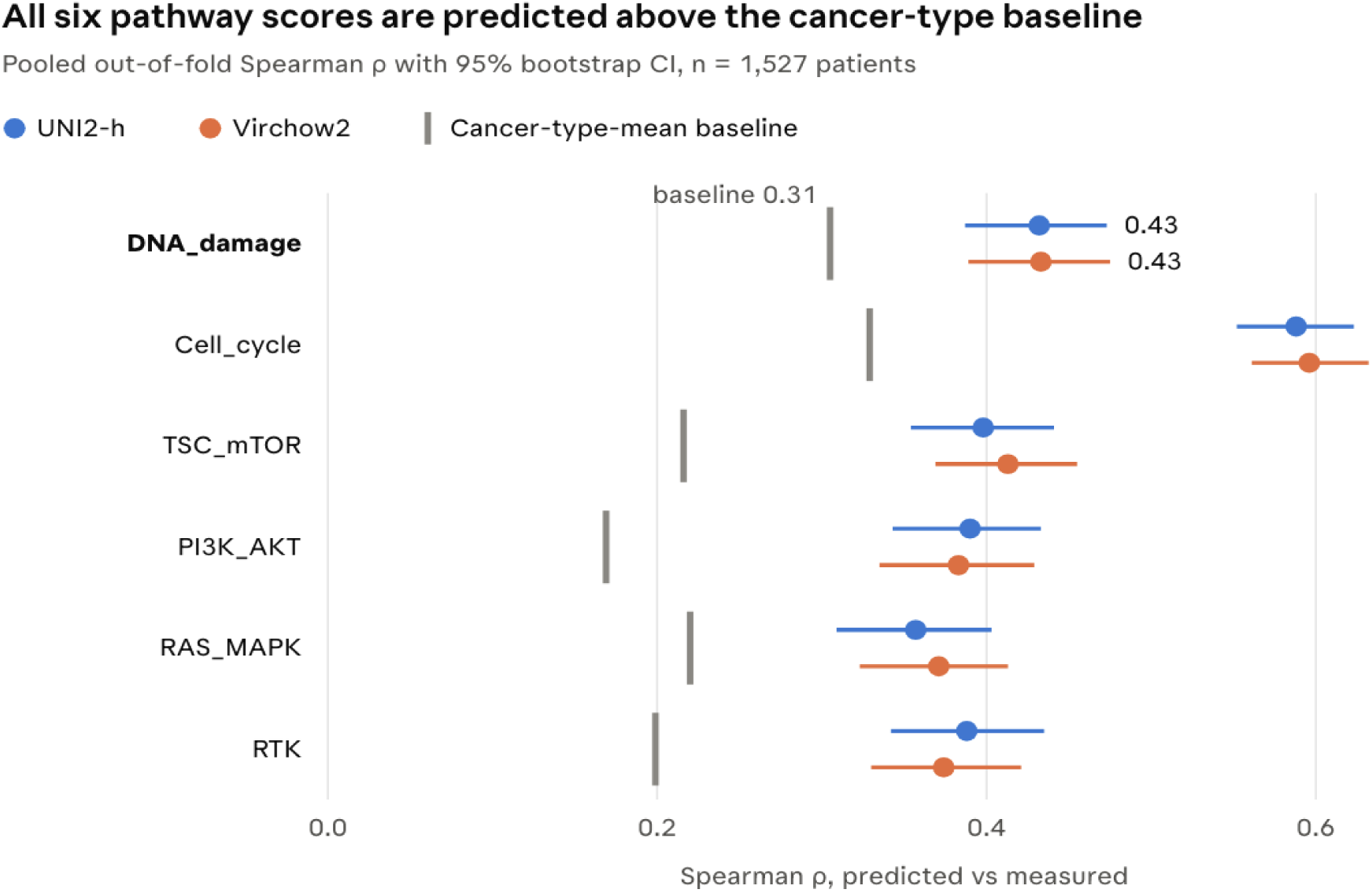
Pooled out-of-fold Spearman ρ (95% bootstrap CI) for each pathway score, with the cancer-type-mean baseline shown as a gray bar.

**Figure 2.**
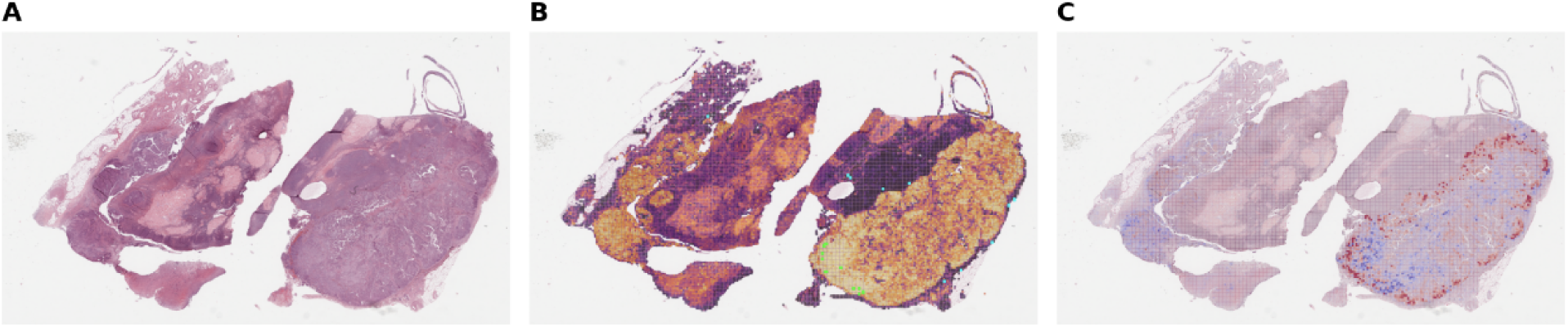
Example held-out ovarian slide, TCGA-25-2399. (A) Raw H&E. (B) UNI2-h ABMIL attention rank map; green marks the top-10 attention tiles and cyan the bottom-10 tiles. (C) UNI2-h Integrated Gradients map for the DNA_damage output; red tiles raise the predicted score and blue tiles lower it.

## Discussion

Routine H&E slides carry a moderate, reproducible signal of DDR protein levels that goes beyond cancer type. The signal was nearly identical with two independently trained foundation models, survived held-out tissue source sites, and disappeared under a within-type permutation. Together these results support the central hypothesis in a narrower form than proposed: H&E predicts the abundance of DDR machinery, and whether it predicts kinase activity itself is still open.

A within-type ρ of about 0.29 means H&E ranks patients’ DNA_damage scores better than chance, but it is far from a patient-level measurement. A baseline that knows only cancer type already reaches ρ = 0.31, most of the pooled ρ of 0.43; the image adds Δρ = +0.13.

Reporting Δρ and within-type ρ alongside pooled ρ was therefore essential; pooled correlation alone would have overstated what the model learned. DeepHRD’s success with a binary genomic label (Bergstrom et al., 2024) suggests that a binarized high-versus low-activity target might be easier to predict.

The attention analysis points to biologically plausible features. High-attention tiles were tumor-dense, with enlarged nuclei and lower chromatin homogeneity, which are classic correlates of genomic instability and aneuploidy (Fernandes, 2024). Lymphocyte-rich stroma received low attention. Agreement between attention and Integrated Gradients (median ρ ≈ 0.70) suggests the attention maps are not arbitrary.

Two caveats limit these conclusions. First, attention and IG come from the same model, so some agreement is built in; it shows internal consistency rather than independent confirmation. Second, tile-quality metrics also differed between high- and low-attention tiles. Some of the morphological contrast may reflect focus or tissue quality rather than tumor biology.

### Limitations

- Cell_cycle was the best-predicted target, and measured DNA_damage and Cell_cycle scores were themselves correlated (Spearman ρ = 0.48 pooled; 0.24–0.50 within cancer types). Part of the DNA_damage signal may reflect proliferation rather than DDR-specific biology.
- 9 of the 11 DNA_damage components are total proteins. The score reflects how much repair machinery is present, not whether ATM and ATR are actively signaling. The proposal’s focus on ATM-CHK2 and ATR-CHK1 activity was not yet tested directly.
- TCGA RPPA was measured on frozen tissue, while the diagnostic slides are FFPE sections from different parts of the tumor. Intratumoral heterogeneity between the two samples adds noise and caps achievable correlation.
- BRCA made up 56% of patients, and OV and UCS together fewer than 8%. Pooled results mostly reflect breast and endometrial cancer, and per-type conclusions for OV and UCS are weak.
- All results come from TCGA cross-validation. Site-grouped folds reduce but do not remove the risk that the model learned TCGA-specific processing artifacts.
- The proposal’s link to platinum and PARP inhibitor response was outside the scope of this term.

### Future directions

1. Train separate models on CHK1_pS345 and CHK2_pT68, the two phosphoproteins that most directly reflect kinase activity.
2. Test whether predicted DNA_damage still correlates with measured DNA_damage after adjusting for the Cell_cycle score, overall and within each cancer type.
3. Add RNA-based HRD and DDR expression signatures as a second ground truth.
4. Validate on an external cohort with matched slides and phosphoproteomics, such as CPTAC breast, endometrial, and ovarian cohorts.
5. Add confounder controls for tumor purity and grade, and compare against another foundation model such as CONCH (Lu et al., 2024).

If these steps confirm a DDR-specific signal, H&E-based scores could eventually help prioritize patients for confirmatory DDR testing. That clinical use would require prospective validation well beyond this study.

### Perspectives

The project ended narrower and more rigorous than the proposal: I set out to predict ATM-CHK2 and ATR-CHK1 kinase activity and finished with a well-controlled prediction of DDR protein abundance, plus a clear plan for reaching kinase activity.

Feedback on my first proposal draft pointed out that five objectives were too many for one term. I restructured the work into two primary aims (cohort assembly and prediction) and one exploratory aim (interpretability), and dropped the treatment-response analysis. I also deferred the proposed cross-tissue transfer test (training on some cancer types and testing on others); the within-type and per-type analyses address part of that question.

I planned to predict CHK1 and CHK2 phosphorylation directly. In practice, I first trained on the six TCGA pathway scores jointly, which gave a stable multi-task setup and let me compare DDR with other pathways. Only after seeing the results did I fully appreciate that the DNA_damage score is mostly total protein. Modeling the phosphoproteins alone is now the first next step.

The proposal specified the original UNI. My first full run used it, and I then moved to UNI2-h and Virchow2. Running two encoders turned out to be one of the strongest checks in the project, because their near-identical results show the signal is not specific to one model’s features.

I originally planned to report MSE, Pearson r, and R^2^ under nested cross-validation. Once I saw that a model predicting only the cancer-type mean already reached ρ ≈ 0.31, I realized pooled correlation alone would be misleading. I added the cancer-type baseline, within-type correlations, site-grouped folds, residualized targets, and a permutation null. These analyses changed how I think about evaluating any model trained on a pooled, multi-type cohort.

### Difficulties

Matching slides to RPPA left 1,527 patients rather than about 1,800, and OV and UCS ended up small. This limited what I could say about individual cancer types. Tiling, normalizing, and embedding 20.4 million tiles with two large encoders required careful Slurm job management and caching of features. Feature extraction ran as four parallel array jobs per encoder, and the stored float32 features came to roughly 125 GB for UNI2-h and 209 GB for Virchow2. Training was cheaper, at about 19 GPU-hours across the eight full cross-validation runs. The most instructive failure was a concurrency bug. Overlapping extraction jobs wrote to the same slide files and left many with valid features but zeroed tile coordinates. This did not affect training, but it would have broken Aim 3. I repaired the coordinates by regenerating them with the same deterministic tiling code and verified them against intact slides before overwriting anything. I then made extraction skip finished slides and made training resume from checkpoints, so that later failed or resubmitted jobs cost time but not results.

Several runs produced similar numbers, and output files did not always record which encoder or fold scheme produced them. At one point I had to reconstruct whether a run used Virchow or Virchow2, and which encoder the robustness run used. Going forward I will write the encoder, fold scheme, and target list into every output file, following JHU Data Services guidance on documenting research data and code (Appendix B).

Additionally, it was tempting to read attention maps as direct evidence of biology.

Finding that tile quality also shifted with attention was a useful reminder to treat these maps as hypotheses.

## Supporting information

RPPA data

GDC_manifest

results

## Appendices

## Appendix A Model and training settings

| **Setting** | **Value** |
|---|---|
| Tile size and resolution | 256x256 px at 0.5 µm/px (~20x) |
| Tissue mask | Otsu threshold on saturation; tiles kept at ≥50% tissue |
| Regression head | Two-layer MLP (512 → 256 → 6), ReLU |
| Stain normalization | Macenko, shared reference |
| Encoders (frozen) | UNI2-h (1,536-d), Virchow2 (2,560-d); UNI (1,024-d) for robustness run |
| Aggregator | Gated attention ABMIL |
| Hidden / attention size | 512 / 256 |
| Dropout | 0.25 |
| Outputs | 6 RPPA pathway scores, trained jointly |
| Loss | Mean squared error, missing targets masked |
| Optimizer | AdamW, learning rate 1x10<sup>-4</sup>, weight decay 1x10<sup>-4</sup>, cosine decay |
| Gradient accumulation | 8 bags per optimizer step |
| Precision | bfloat16 mixed precision |
| Max tiles per bag (training) | 8,000, randomly sampled each step |
| Epochs | Up to 50, early stopping patience 10 |
| Cross-validation | 5 folds by patient, stratified by cancer type, grouped by tissue source site; next fold used for validation |
| Random seed | Base seed 0; fold k trained with seed k; within-type shuffle seed 1000; bootstrap and Aim 3 seed 0. GPU operations not forced to be deterministic. |
| Hardware | Tesla T4 GPU (extraction, training, HoVer-Net, IG); CPU nodes for tiling, export, and statistics |

## Appendix B Data and code documentation

Following JHU Data Services guidance on documenting research data and code, the project repository should include a README, a data dictionary for each output table, and a record of which job produced each result.

**Inputs:** TCGA diagnostic slides (GDC, open access) and the TCGA pan-cancer Level 4 RPPA release (TCGA-RPPA-pancan-clean), downloaded 10/23/2026. Both are public; no protected health information was used.

| Run | Encoder | Folds | Targets | Run folder |
|---|---|---|---|---|
| Initial | UNI | Type-stratified, patient-level (not site-grouped) | 6 pathway scores | 11-2-26/ (train_uni.sbatch) |
| Main | UNI2-h | Type-stratified, site-grouped | 6 pathway scores | uni2h_11-3-26/ (fm_train.sbatch) |
| Main | Virchow2 | Type-stratified, site-grouped | 6 pathway scores | virchow2_11-3-26/ (fm_train.sbatch) |
| Robustness | UNI | Site-held-out | Residualized pathway scores | uni_debias/ (train_debias.sbatch) |
| Null | UNI | Site-held-out | Residualized scores, shuffled within type | uni_debias_null/ (train_debias.sbatch) |
| Null | UNI2-h, Virchow2 | Type-stratified, site-grouped | 6 pathway scores, shuffled within type | uni2h_pathways_null/, virchow2_pathways_null/ (fm_train_null.sbatch) |
| Interpretability | UNI2-h, Virchow2 | Held-out slides (1,527 patients, 1,673 slides) | Attention, IG, HoVer-Net | uni2h_11-3-26/aim2/, virchow2_11-3-26/aim2/ (07_aim2_submit.sh) |

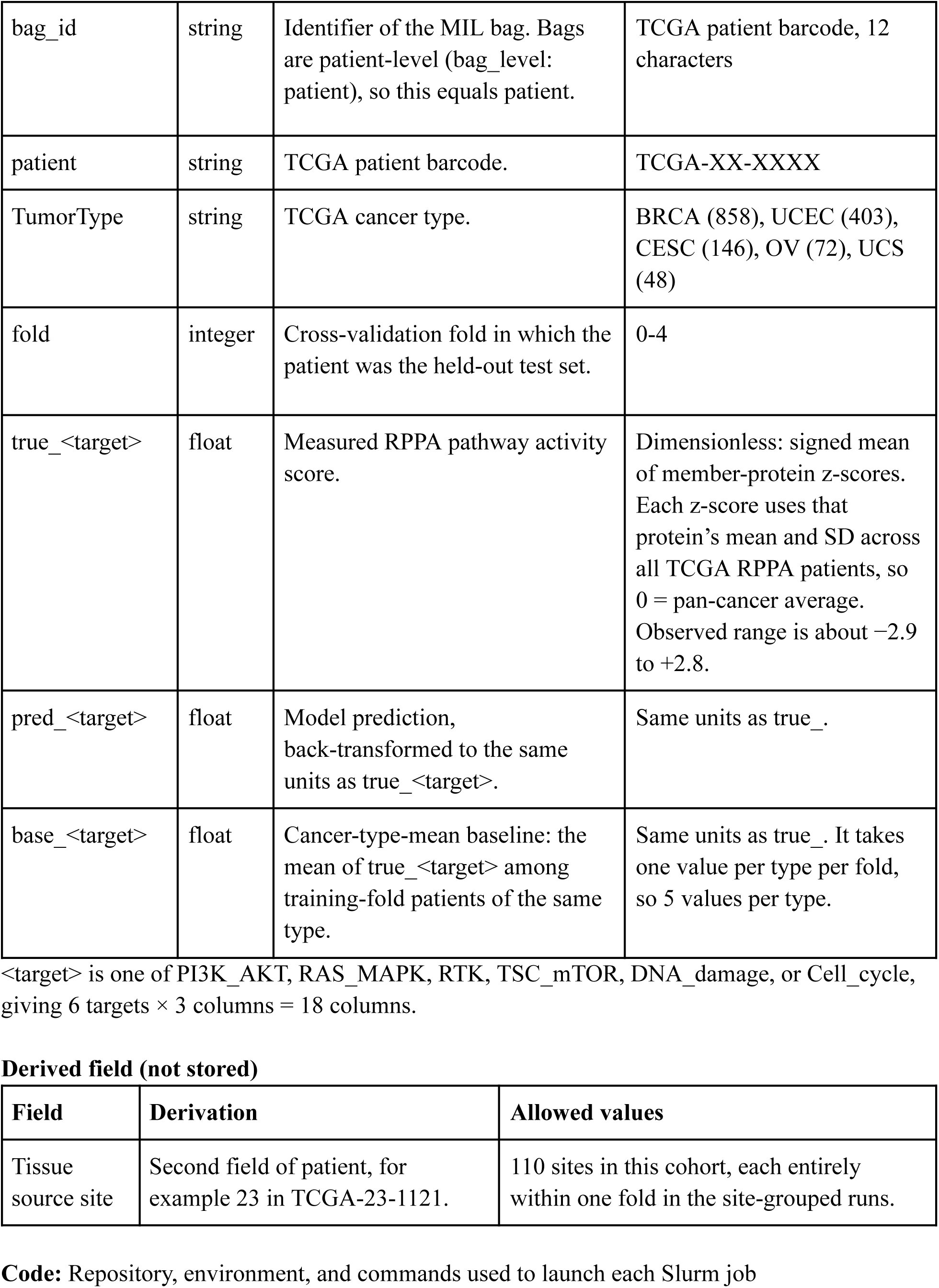

| bag_id | string | Identifier of the MIL bag. Bags are patient-level (bag_level: patient), so this equals patient. | TCGA patient barcode, 12 characters |
|---|---|---|---|
| patient | string | TCGA patient barcode. | TCGA-XX-XXXX |
| TumorType | string | TCGA cancer type. | BRCA (858), UCEC (403), CESC (146), OV (72), UCS (48) |
| fold | integer | Cross-validation fold in which the patient was the held-out test set. | 0-4 |
| true_<target> | float | Measured RPPA pathway activity score. | Dimensionless: signed mean of member-protein z-scores. Each z-score uses that protein's mean and SD across all TCGA RPPA patients, so 0 = pan-cancer average. Observed range is about −2.9 to +2.8. |
| pred_<target> | float | Model prediction, back-transformed to the same units as true_<target>. | Same units as true_. |
| base_<target> | float | Cancer-type-mean baseline: the mean of true_<target> among training-fold patients of the same type. | Same units as true_. It takes one value per type per fold, so 5 values per type. |

<target> is one of PI3K\_AKT, RAS\_MAPK, RTK, TSC\_mTOR, DNA\_damage, or Cell\_cycle, giving 6 targets × 3 columns = 18 columns.
**Derived field (not stored)**
**Field** **Derivation** **Allowed values**
Tissue source site Second field of patient, for example 23 in TCGA-23-1121. 110 sites in this cohort, each entirely within one fold in the site-grouped runs.
**Code:** Repository, environment, and commands used to launch each Slurm job

| **Environment** | **Setup (from the sbatch scripts)** | **Used for** |
|---|---|---|
| pt22_cv (venv) | module load proxy python311 pytorch-py311-cuda12.1-gcc11/2.2.0, then source /projects/wgs_pipeline_dev/envs/pt22_cv/bin/activate. This is Python 3.11, PyTorch 2.2.0, CUDA 12.1. | Download, matching, UNI extraction, QC, all ABMIL training |
| pathfm (conda) | conda activate pathfm, with HF_HUB_OFFLINE=1 set for extraction | UNI2-h and Virchow2 extraction; Aim 3 |
| hovernet (conda) | ~/miniconda3/envs/hovernet/bin/python, with the PanNuke checkpoint in ~/hover_net/checkpoints/ | HoVer-Net nuclear segmentation (Aim 3) |
| **Launch commands:** All jobs run from /projects/wgs_pipeline_dev |  |  |
| **Step** | **Command** | **Output** |
| Download slides | sbatch 01_download_gdc.sh | data/ |
| Match, extract, QC, and train (initial pipeline, ResNet-50) | sbatch run_pipeline.sbatch | wsi_rppa_match/, abmil_data/, abmil_runs/pathways (now 10-28-26) |
| UNI extraction | sbatch --array=0-3 uni_extract.sbatch, then sbatch uni_extract.sbatch to finish and index | abmil_data_uni/ |
| UNI2-h and Virchow2 extraction | sbatch --array=0-3 --export=ALL,ENCODER=uni2h fm_extract.sbatch (same with ENCODER=virchow2) | abmil_data_uni2h/, abmil_data_virchow2/ |
| Coordinate repair | python 07_fix_coords.py --data abmil_data_uni --apply (per data folder, after a dry run) | Repairs the h5 files in place |

| Initial UNI training | sbatch train_uni.sbatch | abmil_runs/uni_pathways (now 11-2-26) |
|---|---|---|
| Robustness | sbatch train_debias.sbatch | abmil_runs/uni_debias |
| Null (UNI) | sbatch --export=ALL,NULL=1 train_debias.sbatch | abmil_runs/uni_debias_null |
| Main runs | sbatch --export=ALL,ENCODER=uni2h fm_train.sbatch (same with ENCODER=virchow2) | abmil_runs/{uni2h,virchow2}_pathways (now *_11-3-26) |
| Null (UNI2-h, Virchow2) | sbatch --export=ALL,ENCODER=uni2h --job-name=fm_train_null fm_train_null.sbatch (same with virchow2) | abmil_runs/{uni2h,virchow2}_pathways_null |
| Aim 3 | bash 07_aim2_submit.sh abmil_runs/uni2h_11-3-26 (same with virchow2_11-3-26). This submits 4 chained jobs: select → heatmaps and tiles → HoVer-Net → features and stats. | <run>/aim2/ |

## Appendix C Full morphometric comparison

| **Feature** | **UNI2-h (SD)** | **q** | **Virchow2 (SD)** | **q** |
|---|---|---|---|---|
| Nuclear density (per mm²) | +0.82 | 1.5e−154 | +0.91 | 5.2e−166 |
| Tumor nuclear solidity | +0.62 | 9.7e−67 | +0.45 | 2.8e−34 |
| Tumor chromatin H mean | −0.37 | 1.4e−18 | −0.24 | 9.5e−8 |
| Tumor nuclear eccentricity | −0.28 | 4.9e−21 | −0.13 | 2.7e−5 |
| Tumor nuclear area CV | +0.26 | 7.4e−23 | +0.31 | 1.3e−24 |
| Tumor chromatin GLCM contrast | +0.23 | 2.6e−31 | +0.38 | 5.1e−43 |

|  |  |  |  |  |
|---|---|---|---|---|
| Lymphocyte density (per mm²) | −0.22 | 2.4e−63 | −0.23 | 8.9e−34 |
| Tumor nuclear circularity | −0.15 | 1.4e−11 | −0.12 | 5.1e−6 |
| Tumor chromatin entropy | −0.03 | 2.8e−9 | +0.08 | 0.72 (n.s.) |
| Tumor chromatin H SD | −0.03 | 0.77 (n.s.) | +0.20 | 1.5e−13 |

Values are the mean per-patient difference (high minus low attention) in feature SD units, tested with a two-sided Wilcoxon signed-rank test with BH correction. Tumor nuclear and chromatin features include patients with tumor nuclei in both tile groups (n = 1,243 UNI2-h; 1,076 Virchow2). For UNI2-h, chromatin entropy is significant despite a small SD difference because the test is rank-based (rank-biserial −0.20).

**Table C2.**
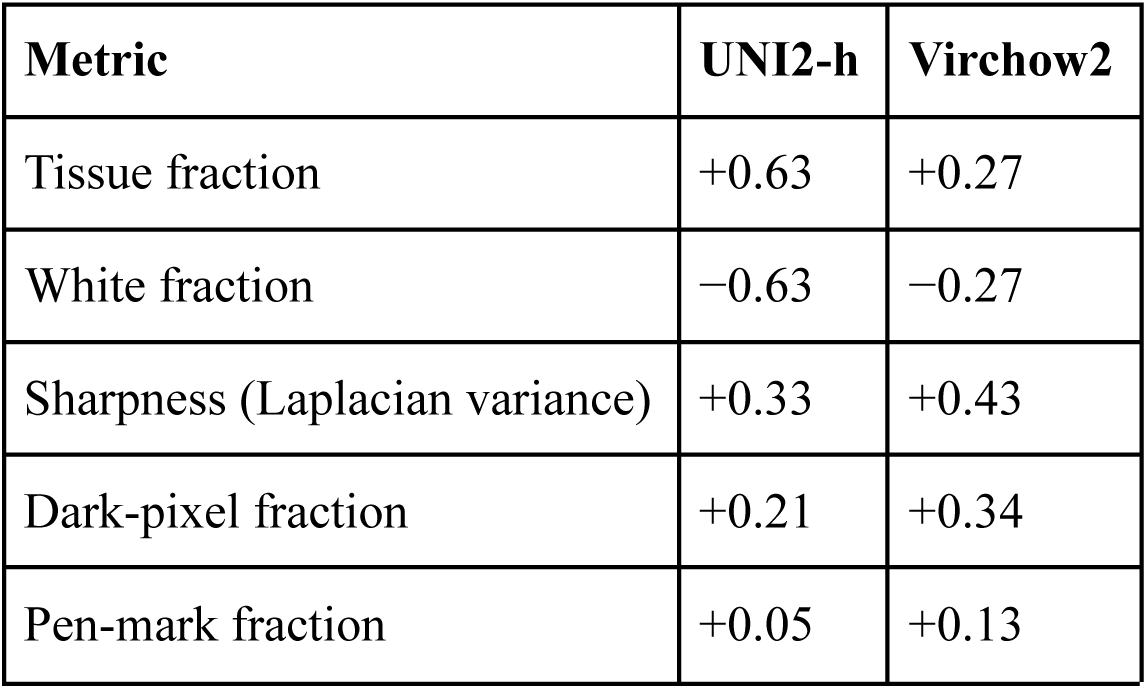
Tile-quality differences, top-10 vs bottom-10 attention tiles (SD units; all q < 1e−17).

| **Metric** | **UNI2-h** | **Virchow2** |
|---|---|---|
| Tissue fraction | +0.63 | +0.27 |
| White fraction | −0.63 | −0.27 |
| Sharpness (Laplacian variance) | +0.33 | +0.43 |
| Dark-pixel fraction | +0.21 | +0.34 |
| Pen-mark fraction | +0.05 | +0.13 |

## Appendix D DNA_damage score components

| **Group** | **Proteins** | **Type** |
|---|---|---|
| Damage sensing | ATM, MRE11, RAD50 | Total protein |
| Homologous recombination | BRCA2, RAD51 | Total protein |
| Non-homologous end joining | KU80, 53BP1 | Total protein |
| Other | P53, XRCC1 | Total protein |
| Checkpoint kinase activity | CHK1_pS345 (ATR target), CHK2_pT68 (ATM target) | Phosphoprotein |

