## Supplementary material for "Deep Learning Prediction of DNA Damage Response Protein Signaling from H&E Histopathology and Reverse-Phase Protein Arrays in Pan-Gynecologic Cancers": results

### Predicting RPPA kinase-signalling pathway activity from H&E whole-slide images in pan-gynecologic cancers

Results package for manuscript submission: attention-based multiple-instance learning on pathology foundation-model features

Samantha Lane · M.S. Bioinformatics thesis, Johns Hopkins University · generated November 16, 2026

#### Key results

- Cohort: 1,527 TCGA patients across five gynecologic and breast cancers (BRCA, UCEC, CESC, OV, UCS), 1,673 FFPE diagnostic slides, each matched to TCGA RPPA proteomics; six kinase-signalling pathway scores predicted jointly.
- Primary model (UNI2-h features + ABMIL), pooled out-of-fold Spearman ρ ranged 0.36 to 0.59 across pathways; highest for Cell cycle (ρ = 0.588) and DNA damage (ρ = 0.432).
- The model beat a cancer-type-mean baseline for 6/6 pathways (Δρ 95% CI above 0; Δρ 0.13 to 0.26), and kept signal within cancer type for 6/6 pathways (within-type ρ 0.28 to 0.46).
- Permutation control (targets shuffled within cancer type): within-type ρ fell to -0.07 to 0.03 and Δρ to -0.04 to -0.01, so the real-label signal is not explained by tissue type alone.
- Replication with Virchow2 features gave within-type ρ 0.29 to 0.48 (largest per-pathway difference from UNI2-h: 0.032).
- With folds stratified by tissue-source site and type-residualized targets (UNI features), within-type ρ stayed at 0.26 to 0.42.
- Aim 2 (DNA damage model): high- vs low-attention tiles differed on 13/14 HoVer-Net nuclear/microenvironment features (BH $q < 0.05$); attention agreed with integrated gradients (per-fold median Spearman 0.72); 6 pathomic tile clusters, all enriched by attention group.

#### Contents


- Cohort and methods summary
- Table 1. Pooled out-of-fold performance (UNI2-h primary, Virchow2 replication)
- Figure 1. Predicted vs measured pathway scores
- Figure 2. Signal beyond cancer type: model vs permutation null and site-stratified CV
- Table 2. Encoder and control comparison
- Figure 3 / Table S1. Performance within each cancer type
- Table S2 / Figure S1. Fold-level consistency and training curves
- Aim 2: interpretability (attention vs HoVer-Net features, IG agreement, pathomic clusters, heatmaps)
- Limitations, items still needed for submission, and reproducibility record

### Cohort and methods summary

Values on this page are read from the run configs and matched-cohort files; check wording against the manuscript methods before submission.

##### Analysed cohort (patients with ≥1 FFPE DX slide and RPPA)

| Cancer type | Patients | DX slides | % of patients | DNA damage score mean (SD) | Cell cycle score mean (SD) |
|---|---|---|---|---|---|
| BRCA | 858 | 917 | 56.2 | -0.16 (0.35) | -0.17 (0.55) |
| UCEC | 403 | 451 | 26.4 | +0.08 (0.37) | +0.12 (0.51) |
| CESC | 146 | 152 | 9.6 | +0.02 (0.38) | +0.44 (0.45) |
| OV | 72 | 73 | 4.7 | +0.17 (0.35) | +0.12 (0.42) |
| UCS | 48 | 80 | 3.1 | +0.64 (0.39) | +0.47 (0.53) |
| Total | 1527 | 1673 | 100.0 | -0.04 (0.40) | -0.00 (0.57) |

#### Data

- TCGA BRCA, CESC, OV, UCEC and UCS diagnostic FFPE (DX) whole-slide images from GDC, matched at the patient level to the TCGA pan-cancer RPPA level-4 matrix (198 antibodies). Frozen-section slides excluded.
- Targets: six kinase-signalling pathway scores (PI3K/AKT, RAS/MAPK, RTK, TSC/mTOR, DNA damage response, cell cycle), each the signed mean of member-protein z-scores (z computed over all pan-cancer RPPA patients).

#### Image processing and model

- Tissue tiles of 256 px at 0.5 µm/px (~20×), tissue fraction ≥ 0.5, Macenko stain normalisation; tile embeddings from frozen foundation models: UNI2-h (1536-d), Virchow2 (2560-d), UNI (v1) (1024-d).
- Attention-based MIL (ABMIL) multi-output regression with one bag per patient (all slides pooled); hidden 512, attention dim 256, dropout 0.25; lr 0.0001, weight decay 0.0001, gradient accumulation 8; ≤50 epochs with early stopping (patience 10); ≤8000 tiles sampled per training bag; targets z-scored with training-fold mean/SD; seed 0.

#### Evaluation

- 5-fold patient-level cross-validation with a fixed fold file shared across encoders; all headline metrics computed on pooled out-of-fold predictions for every patient.
- 95% CIs from 1000 patient bootstrap resamples; Benjamini–Hochberg q across pathways. Baseline = training-fold cancer-type mean (what a model gets by recognising tissue type). $\Delta\rho$ = model $\rho$ − baseline $\rho$. Within-type $\rho$ = Spearman correlation after removing cancer-type differences.
- Controls: (i) permutation null, targets shuffled within cancer type (keeps type signal, breaks the patient-level link); (ii) folds stratified by TCGA tissue-source site with type-residualised targets (UNI features) to test for site/batch leakage.
- Aim 2 interpretability: per held-out slide, top/bottom-K attention tiles; HoVer-Net (PanNuke) nuclear segmentation and morphometry; patient-level Wilcoxon tests; integrated gradients (IG); k-means pathomic clusters with k chosen by silhouette.

### Table 1. Pooled out-of-fold performance

n = 1,527 patients, 5-fold CV. Upper: UNI2-h (primary). Lower: Virchow2 (replication, same folds). Main claims should rest on within-type ρ and Δρ, not raw ρ, because pathway scores differ strongly between cancer types.

#### 1a. UNI2-h + ABMIL

| Pathway | Spearman ρ [95% CI] | Pearson r | $R^2$ [95% CI] | Within-type ρ [95% CI] | Type-mean baseline ρ | Δρ vs baseline [95% CI] | q (ρ) | q (within) |
|---|---|---|---|---|---|---|---|---|
| PI3K/AKT | 0.390 [0.34, 0.43] | 0.410 | 0.167 [0.13, 0.20] | 0.319 [0.27, 0.36] | 0.169 | 0.221 [0.16, 0.28] | 1.9e-56 | 5.2e-37 |
| RAS/MAPK | 0.357 [0.31, 0.40] | 0.353 | 0.118 [0.08, 0.16] | 0.277 [0.23, 0.32] | 0.220 | 0.138 [0.08, 0.19] | 3.6e-47 | 2.6e-28 |
| RTK | 0.388 [0.34, 0.43] | 0.376 | 0.138 [0.10, 0.17] | 0.320 [0.27, 0.37] | 0.199 | 0.188 [0.13, 0.24] | 7.6e-56 | 4.8e-37 |
| TSC/mTOR | 0.398 [0.35, 0.44] | 0.409 | 0.158 [0.11, 0.20] | 0.287 [0.24, 0.33] | 0.216 | 0.182 [0.12, 0.24] | 6.5e-59 | 3.1e-30 |
| DNA damage | 0.432 [0.39, 0.47] | 0.477 | 0.226 [0.18, 0.27] | 0.294 [0.25, 0.34] | 0.305 | 0.127 [0.09, 0.16] | 5.9e-70 | 1.7e-31 |
| Cell cycle | 0.588 [0.55, 0.62] | 0.561 | 0.309 [0.26, 0.35] | 0.462 [0.42, 0.50] | 0.329 | 0.259 [0.22, 0.30] | 9.4e-142 | 8.6e-81 |

#### 1b. Virchow2 + ABMIL

| Pathway | Spearman ρ [95% CI] | Pearson r | $R^2$ [95% CI] | Within-type ρ [95% CI] | Type-mean baseline ρ | Δρ vs baseline [95% CI] | q (ρ) | q (within) |
|---|---|---|---|---|---|---|---|---|
| PI3K/AKT | 0.383 [0.34, 0.43] | 0.411 | 0.165 [0.12, 0.20] | 0.332 [0.28, 0.38] | 0.169 | 0.214 [0.16, 0.27] | 2.8e-54 | 5.3e-40 |
| RAS/MAPK | 0.371 [0.32, 0.41] | 0.362 | 0.125 [0.08, 0.16] | 0.290 [0.24, 0.34] | 0.220 | 0.152 [0.10, 0.21] | 4.4e-51 | 8.1e-31 |
| RTK | 0.374 [0.33, 0.42] | 0.367 | 0.132 [0.10, 0.17] | 0.288 [0.24, 0.34] | 0.199 | 0.175 [0.12, 0.23] | 7.2e-52 | 1.8e-30 |
| TSC/mTOR | 0.413 [0.37, 0.46] | 0.415 | 0.166 [0.12, 0.20] | 0.301 [0.25, 0.35] | 0.216 | 0.197 [0.15, 0.25] | 1.0e-63 | 4.7e-33 |
| DNA damage | 0.433 [0.39, 0.48] | 0.482 | 0.228 [0.18, 0.27] | 0.303 [0.26, 0.35] | 0.305 | 0.128 [0.09, 0.16] | 3.1e-70 | 1.7e-33 |
| Cell cycle | 0.596 [0.56, 0.63] | 0.568 | 0.319 [0.27, 0.36] | 0.478 [0.43, 0.52] | 0.329 | 0.268 [0.22, 0.32] | 6.4e-147 | 3.6e-87 |

ρ, Spearman correlation between predicted and measured score; $R^2$, coefficient of determination on the original scale; within-type ρ, correlation after removing cancer-type means; Δρ, improvement over the cancer-type-mean baseline. q, Benjamini–Hochberg adjusted. p/q are approximate because pooled CV predictions are not fully independent; CIs are patient-bootstrap percentiles.

### Figure 1. Predicted vs measured pathway activity (UNI2-h, out-of-fold)

Each point is one held-out patient, coloured by cancer type. Line: least-squares fit. ρ: pooled Spearman; ρw: within-type Spearman.

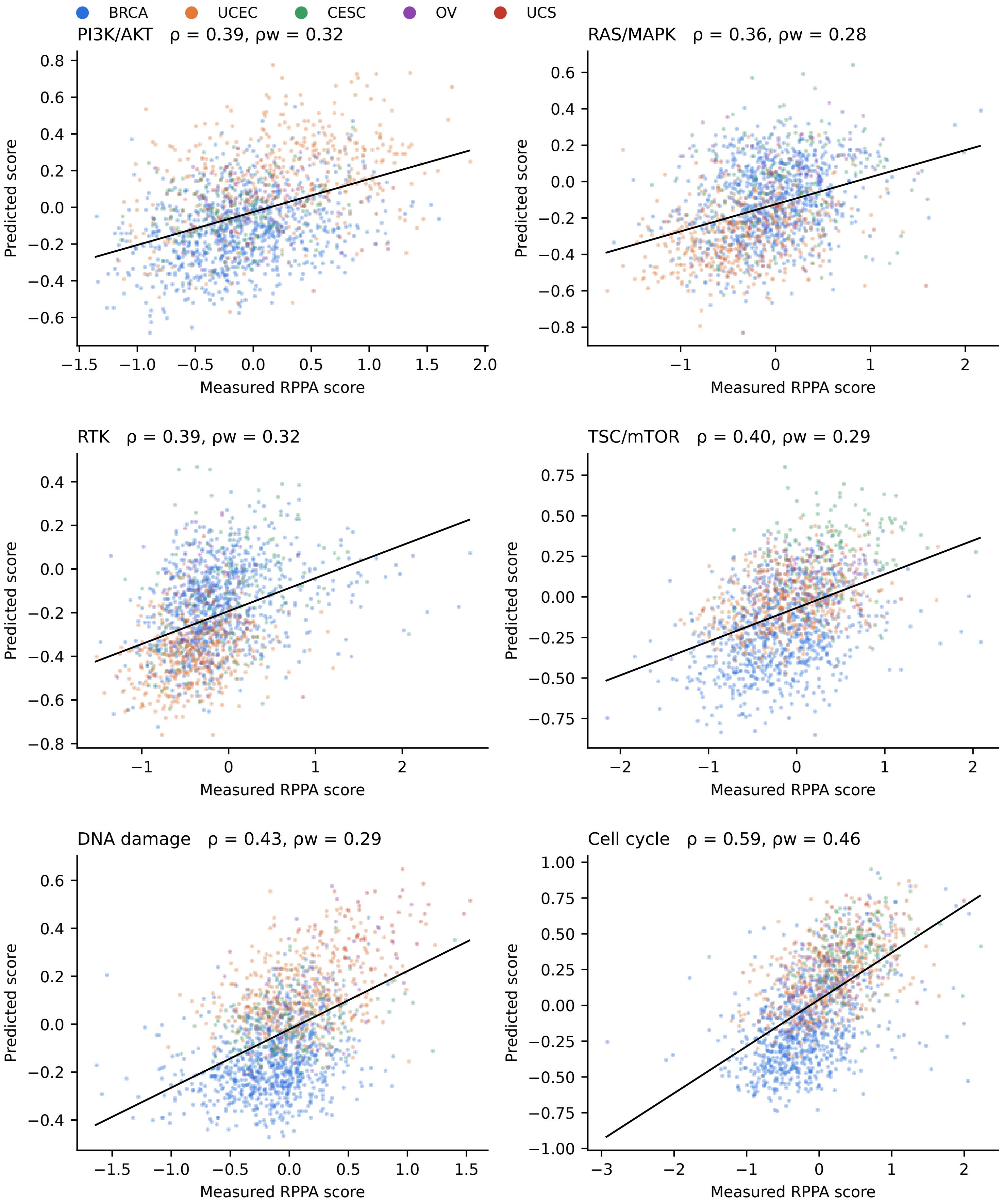

### Figure 2. Signal beyond cancer type

Pooled estimates with 95% bootstrap CIs. Left: within-cancer-type Spearman ρ. Right: Δρ over the cancer-type-mean baseline. Grey/pale points are permutation nulls (targets shuffled within cancer type); they should sit at or below zero.

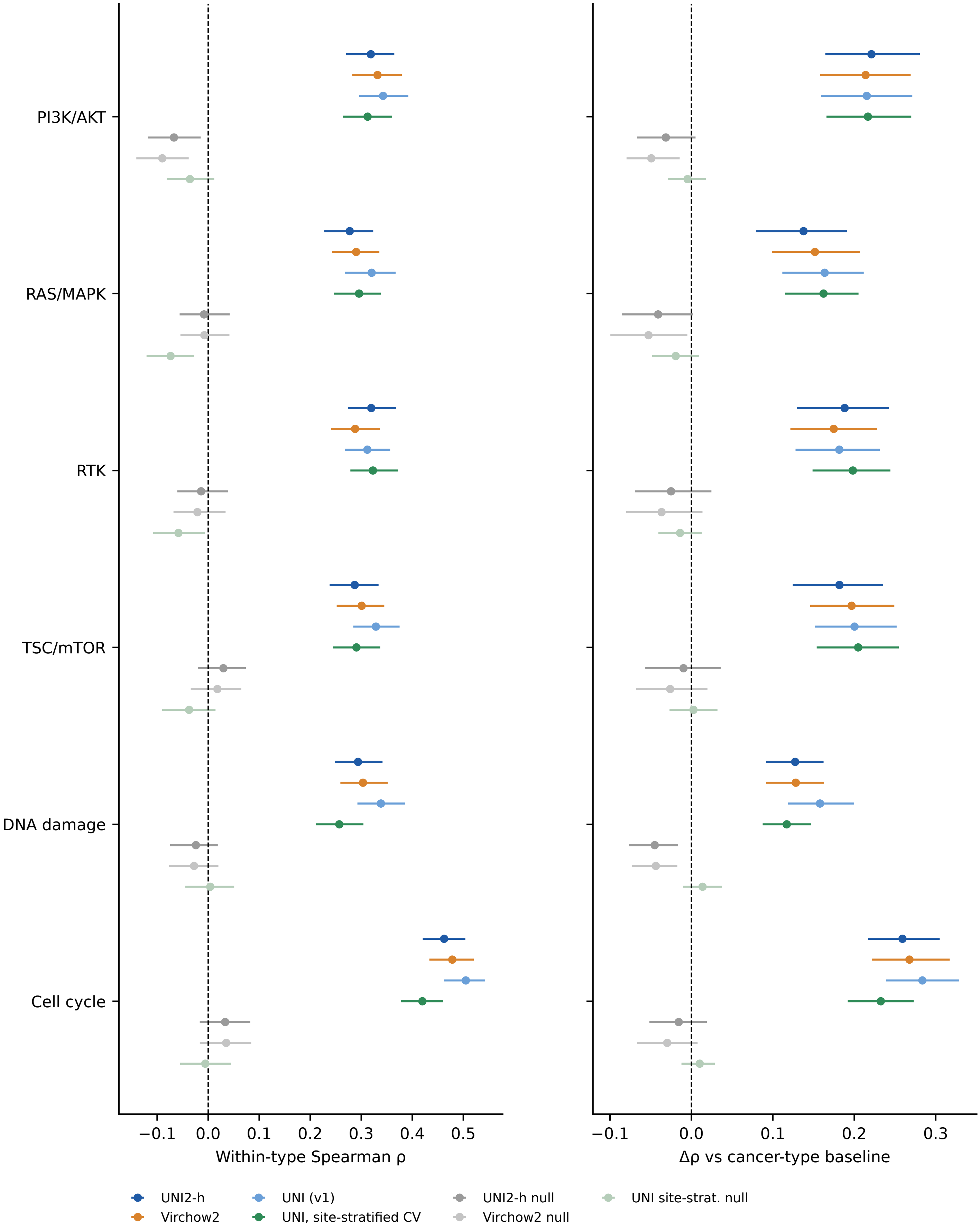

### Table 2. Encoder and control comparison

Pooled out-of-fold estimates [95% CI] for every training run. Null rows shuffle targets within cancer type. Site-stratified rows use UNI features, folds stratified by tissue-source site and type-residualised targets.

#### 2a. Within-cancer-type Spearman ρ

| Model / control | PI3K/AKT | RAS/MAPK | RTK | TSC/mTOR | DNA damage | Cell cycle |
|---|---|---|---|---|---|---|
| UNI2-h (n=1,527) | 0.319 [0.27, 0.36] | 0.277 [0.23, 0.32] | 0.320 [0.27, 0.37] | 0.287 [0.24, 0.33] | 0.294 [0.25, 0.34] | 0.462 [0.42, 0.50] |
| Virchow2 (n=1,527) | 0.332 [0.28, 0.38] | 0.290 [0.24, 0.34] | 0.288 [0.24, 0.34] | 0.301 [0.25, 0.35] | 0.303 [0.26, 0.35] | 0.478 [0.43, 0.52] |
| UNI (v1) (n=1,527) | 0.343 [0.30, 0.39] | 0.320 [0.27, 0.37] | 0.312 [0.27, 0.36] | 0.329 [0.28, 0.38] | 0.339 [0.29, 0.39] | 0.505 [0.46, 0.54] |
| UNI, site-stratified CV (n=1,527) | 0.312 [0.26, 0.36] | 0.296 [0.25, 0.34] | 0.323 [0.28, 0.37] | 0.291 [0.24, 0.34] | 0.257 [0.21, 0.30] | 0.420 [0.38, 0.46] |
| UNI2-h null (n=1,527) | -0.067 [-0.12, -0.01] | -0.008 [-0.06, 0.04] | -0.014 [-0.06, 0.04] | 0.030 [-0.02, 0.07] | -0.024 [-0.07, 0.02] | 0.033 [-0.02, 0.08] |
| Virchow2 null (n=1,527) | -0.090 [-0.14, -0.04] | -0.008 [-0.05, 0.04] | -0.021 [-0.07, 0.03] | 0.018 [-0.03, 0.06] | -0.028 [-0.08, 0.02] | 0.035 [-0.02, 0.08] |
| UNI site-strat. null (n=1,527) | -0.036 [-0.08, 0.01] | -0.074 [-0.12, -0.03] | -0.058 [-0.11, -0.01] | -0.037 [-0.09, 0.01] | 0.004 [-0.04, 0.05] | -0.006 [-0.06, 0.04] |

#### 2b. Δρ over the cancer-type-mean baseline

| Model / control | PI3K/AKT | RAS/MAPK | RTK | TSC/mTOR | DNA damage | Cell cycle |
|---|---|---|---|---|---|---|
| UNI2-h (n=1,527) | 0.221 [0.16, 0.28] | 0.138 [0.08, 0.19] | 0.188 [0.13, 0.24] | 0.182 [0.12, 0.24] | 0.127 [0.09, 0.16] | 0.259 [0.22, 0.30] |
| Virchow2 (n=1,527) | 0.214 [0.16, 0.27] | 0.152 [0.10, 0.21] | 0.175 [0.12, 0.23] | 0.197 [0.15, 0.25] | 0.128 [0.09, 0.16] | 0.268 [0.22, 0.32] |
| UNI (v1) (n=1,527) | 0.215 [0.16, 0.27] | 0.164 [0.11, 0.21] | 0.182 [0.13, 0.23] | 0.200 [0.15, 0.25] | 0.158 [0.12, 0.20] | 0.284 [0.24, 0.33] |
| UNI, site-stratified CV (n=1,527) | 0.217 [0.17, 0.27] | 0.162 [0.12, 0.21] | 0.198 [0.15, 0.24] | 0.205 [0.15, 0.25] | 0.117 [0.09, 0.15] | 0.233 [0.19, 0.27] |
| UNI2-h null (n=1,527) | -0.031 [-0.07, 0.01] | -0.041 [-0.09, 0.00] | -0.025 [-0.07, 0.02] | -0.010 [-0.06, 0.04] | -0.045 [-0.08, -0.02] | -0.016 [-0.05, 0.02] |
| Virchow2 null (n=1,527) | -0.049 [-0.08, -0.01] | -0.053 [-0.10, -0.00] | -0.037 [-0.08, 0.01] | -0.026 [-0.07, 0.02] | -0.044 [-0.07, -0.02] | -0.030 [-0.07, 0.01] |
| UNI site-strat. null (n=1,527) | -0.005 [-0.03, 0.02] | -0.019 [-0.05, 0.01] | -0.014 [-0.04, 0.01] | 0.003 [-0.03, 0.03] | 0.014 [-0.01, 0.04] | 0.010 [-0.01, 0.03] |

Shaded rows: permutation nulls. UNI (v1) is an earlier run with 1,024-d UNI features whose config records default folds (abmil_data_uni/bags.csv) rather than the fold file used for UNI2-h and Virchow2; confirm the fold assignment matches before presenting it side by side.

### Figure 3. Performance within each cancer type

Spearman ρ between predicted and measured scores computed separately in each cancer type (pooled out-of-fold). * BH q < 0.05. Patients per type shown in column labels.

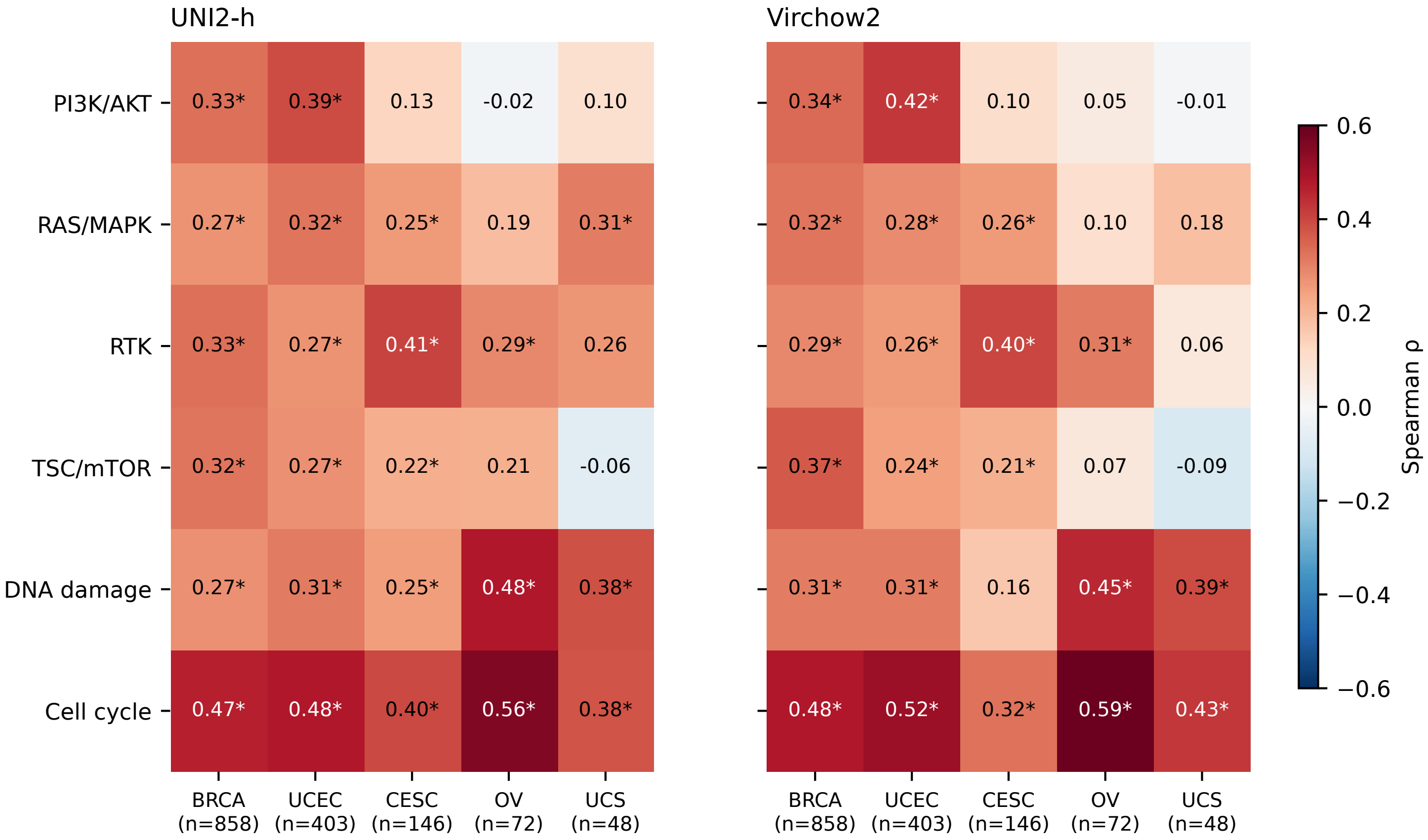


#### Reading the per-type results

- BRCA and UCEC carry most patients and give the tightest intervals; OV (n≈72) and UCS (n≈48) have wide CIs, so per-type claims there should be framed as exploratory.
- Full per-type estimates with 95% CIs and q-values for both encoders are in Table S1 (next page) and tables/tableS1_within_cancer_type.csv.

### Table S1. Spearman ρ within each cancer type, both encoders

Pooled out-of-fold predictions split by cancer type; 95% bootstrap CIs; BH q across pathway × type tests within each encoder.

| Pathway | Type | n | UNI2-h ρ [95% CI] | UNI2-h q | Virchow2 ρ [95% CI] | Virchow2 q |
|---|---|---|---|---|---|---|
| PI3K/AKT | BRCA | 858 | 0.329 [0.27, 0.38] | 3.0e-22 | 0.342 [0.28, 0.40] | 5.0e-24 |
| PI3K/AKT | UCEC | 403 | 0.392 [0.30, 0.48] | 1.4e-15 | 0.423 [0.34, 0.50] | 2.5e-18 |
| PI3K/AKT | CESC | 146 | 0.127 [-0.04, 0.29] | 0.140 | 0.104 [-0.06, 0.27] | 0.274 |
| PI3K/AKT | OV | 72 | -0.018 [-0.27, 0.24] | 0.878 | 0.053 [-0.19, 0.30] | 0.703 |
| PI3K/AKT | UCS | 48 | 0.098 [-0.21, 0.37] | 0.542 | -0.006 [-0.31, 0.28] | 0.967 |
| RAS/MAPK | BRCA | 858 | 0.272 [0.21, 0.33] | 2.0e-15 | 0.319 [0.26, 0.38] | 5.7e-21 |
| RAS/MAPK | UCEC | 403 | 0.323 [0.23, 0.41] | 9.7e-11 | 0.283 [0.19, 0.37] | 2.2e-08 |
| RAS/MAPK | CESC | 146 | 0.254 [0.09, 0.39] | 0.003 | 0.257 [0.09, 0.39] | 0.003 |
| RAS/MAPK | OV | 72 | 0.187 [-0.07, 0.42] | 0.133 | 0.100 [-0.16, 0.34] | 0.482 |
| RAS/MAPK | UCS | 48 | 0.309 [-0.01, 0.59] | 0.043 | 0.181 [-0.13, 0.46] | 0.274 |
| RTK | BRCA | 858 | 0.329 [0.27, 0.40] | 3.0e-22 | 0.288 [0.22, 0.36] | 2.8e-17 |
| RTK | UCEC | 403 | 0.268 [0.17, 0.35] | 1.2e-07 | 0.257 [0.16, 0.34] | 4.4e-07 |
| RTK | CESC | 146 | 0.407 [0.26, 0.54] | 7.7e-07 | 0.401 [0.26, 0.53] | 1.2e-06 |
| RTK | OV | 72 | 0.288 [0.07, 0.46] | 0.019 | 0.311 [0.11, 0.50] | 0.012 |
| RTK | UCS | 48 | 0.265 [-0.02, 0.55] | 0.086 | 0.061 [-0.27, 0.39] | 0.703 |
| TSC/mTOR | BRCA | 858 | 0.321 [0.26, 0.37] | 2.8e-21 | 0.370 [0.31, 0.43] | 5.6e-28 |
| TSC/mTOR | UCEC | 403 | 0.274 [0.18, 0.36] | 6.3e-08 | 0.244 [0.15, 0.34] | 1.5e-06 |
| TSC/mTOR | CESC | 146 | 0.220 [0.07, 0.37] | 0.011 | 0.211 [0.05, 0.36] | 0.015 |
| TSC/mTOR | OV | 72 | 0.213 [-0.03, 0.42] | 0.086 | 0.068 [-0.17, 0.29] | 0.632 |
| TSC/mTOR | UCS | 48 | -0.064 [-0.38, 0.25] | 0.691 | -0.094 [-0.41, 0.25] | 0.608 |
| DNA damage | BRCA | 858 | 0.272 [0.21, 0.33] | 2.0e-15 | 0.306 [0.25, 0.36] | 2.4e-19 |
| DNA damage | UCEC | 403 | 0.310 [0.22, 0.40] | 6.4e-10 | 0.307 [0.22, 0.40] | 1.0e-09 |
| DNA damage | CESC | 146 | 0.251 [0.09, 0.39] | 0.004 | 0.164 [-0.00, 0.32] | 0.066 |
| DNA damage | OV | 72 | 0.480 [0.27, 0.64] | 3.7e-05 | 0.452 [0.25, 0.61] | 1.3e-04 |
| DNA damage | UCS | 48 | 0.383 [0.12, 0.58] | 0.011 | 0.391 [0.16, 0.58] | 0.009 |
| Cell cycle | BRCA | 858 | 0.467 [0.41, 0.52] | 3.4e-46 | 0.478 [0.42, 0.54] | 8.8e-49 |
| Cell cycle | UCEC | 403 | 0.482 [0.40, 0.56] | 1.2e-23 | 0.515 [0.44, 0.59] | 1.1e-27 |
| Cell cycle | CESC | 146 | 0.397 [0.25, 0.53] | 1.4e-06 | 0.324 [0.16, 0.47] | 1.3e-04 |
| Cell cycle | OV | 72 | 0.558 [0.36, 0.71] | 7.7e-07 | 0.589 [0.39, 0.74] | 1.4e-07 |
| Cell cycle | UCS | 48 | 0.377 [0.08, 0.62] | 0.012 | 0.426 [0.14, 0.64] | 0.004 |

### Table S2 and Figure S1. Fold-level consistency and training curves

Per-fold within-type ρ and Δρ on each held-out fold (folds 0–4). Consistent positive signs across folds support that the pooled signal is not driven by one split.

| Encoder | Pathway | Within-type ρ by fold | Folds >0 | Δρ by fold | Folds Δ>0 |
|---|---|---|---|---|---|
| UNI2-h | PI3K/AKT | 0.31 0.36 0.26 0.35 0.37 | 5/5 | 0.14 0.25 0.16 0.07 0.15 | 5/5 |
| UNI2-h | RAS/MAPK | 0.35 0.33 0.17 0.33 0.27 | 5/5 | 0.13 0.18 0.01 0.08 0.10 | 5/5 |
| UNI2-h | RTK | 0.40 0.35 0.23 0.33 0.29 | 5/5 | 0.19 0.20 0.04 0.12 0.11 | 5/5 |
| UNI2-h | TSC/mTOR | 0.30 0.31 0.30 0.33 0.26 | 5/5 | 0.08 0.21 0.20 0.11 0.08 | 5/5 |
| UNI2-h | DNA damage | 0.37 0.24 0.26 0.31 0.31 | 5/5 | 0.11 0.08 0.06 0.13 0.10 | 5/5 |
| UNI2-h | Cell cycle | 0.53 0.45 0.52 0.56 0.37 | 5/5 | 0.20 0.20 0.21 0.28 0.16 | 5/5 |
| Virchow2 | PI3K/AKT | 0.27 0.41 0.31 0.35 0.44 | 5/5 | 0.11 0.25 0.18 0.00 0.20 | 5/5 |
| Virchow2 | RAS/MAPK | 0.32 0.35 0.21 0.33 0.18 | 5/5 | 0.10 0.19 0.08 0.09 0.05 | 5/5 |
| Virchow2 | RTK | 0.38 0.31 0.30 0.34 0.14 | 5/5 | 0.18 0.17 0.12 0.13 0.02 | 5/5 |
| Virchow2 | TSC/mTOR | 0.22 0.32 0.38 0.30 0.29 | 5/5 | 0.03 0.18 0.30 0.10 0.11 | 5/5 |
| Virchow2 | DNA damage | 0.35 0.28 0.35 0.32 0.31 | 5/5 | 0.09 0.06 0.10 0.13 0.08 | 5/5 |
| Virchow2 | Cell cycle | 0.54 0.51 0.47 0.58 0.34 | 5/5 | 0.21 0.23 0.18 0.30 0.13 | 5/5 |

Figure S1a. UNI2-h; best epochs {0: 6, 1: 5, 2: 5, 3: 7, 4: 3}

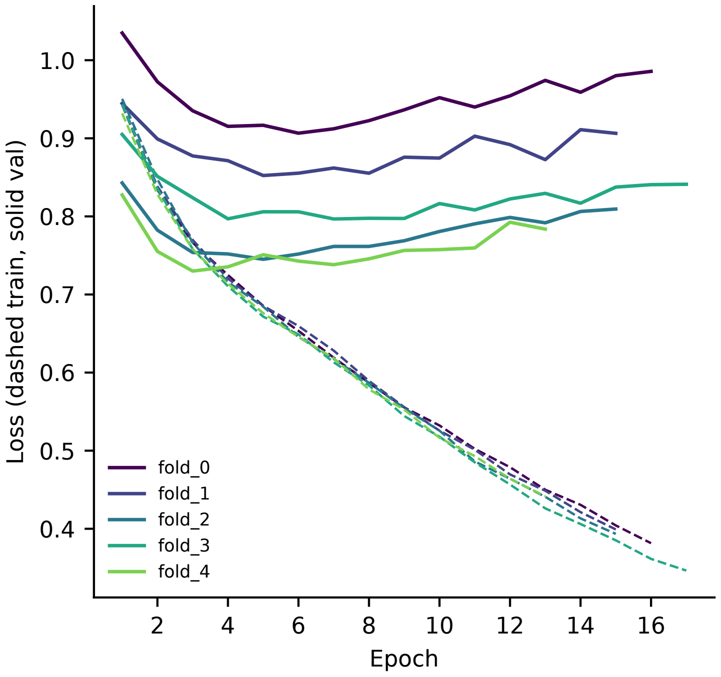


Figure S1b. Virchow2; best epochs {0: 7, 1: 8, 2: 4, 3: 7, 4: 3}

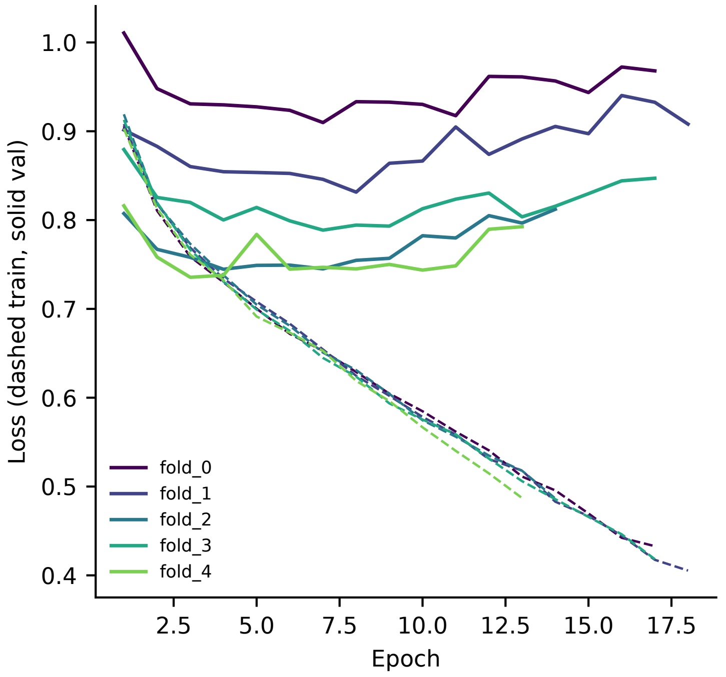

### Table 3. Aim 2: what high-attention tiles look like (DNA damage model)

UNI2-h model; for each of 1,527 held-out patients, the top-10 and bottom-10 attention tiles per slide were segmented with HoVer-Net. Paired patient-level Wilcoxon tests of high minus low; Virchow2 column shows the replication.

| Family | Feature | Patients | Median high | Median low | Std. diff | % pts high>low | Rank-biserial | q (BH) | Virchow2 r_rb (q) |
|---|---|---|---|---|---|---|---|---|---|
| nuclear/TME | tumor_nuc_area_median_um2 | 1243 | 39.7 | 25.1 | +1.26 | 95 | +0.97 | 5.6e-190 | +0.89 (6.3e-142) |
| nuclear/TME | tumor_nuc_area_cv | 1243 | 0.512 | 0.474 | +0.26 | 65 | +0.32 | 7.4e-23 | +0.36 (1.3e-24) |
| nuclear/TME | tumor_nuc_circularity_median | 1243 | 0.842 | 0.852 | -0.15 | 39 | -0.22 | 1.4e-11 | -0.16 (5.1e-06) |
| nuclear/TME | tumor_nuc_solidity_median | 1243 | 0.959 | 0.955 | +0.62 | 70 | +0.57 | 9.7e-67 | +0.43 (2.8e-34) |
| nuclear/TME | tumor_nuc_eccentricity_median | 1243 | 0.708 | 0.723 | -0.28 | 38 | -0.31 | 4.9e-21 | -0.15 (2.7e-05) |
| nuclear/TME | tumor_chromatin_H_mean | 1243 | 0.0601 | 0.0644 | -0.37 | 40 | -0.29 | 1.4e-18 | -0.19 (9.5e-08) |
| nuclear/TME | tumor_chromatin_H_sd | 1243 | 0.019 | 0.019 | -0.03 | 50 | -0.01 | 0.767 | +0.26 (1.5e-13) |
| nuclear/TME | tumor_chromatin_glcm_contrast | 1243 | 2.06 | 1.83 | +0.23 | 64 | +0.38 | 2.6e-31 | +0.49 (5.1e-43) |
| nuclear/TME | tumor_chromatin_glcm_homogeneity | 1243 | 0.574 | 0.606 | -0.42 | 30 | -0.54 | 1.2e-60 | -0.61 (4.6e-66) |
| nuclear/TME | tumor_chromatin_entropy | 1243 | 3.26 | 3.29 | -0.03 | 40 | -0.20 | 2.8e-09 | +0.01 (0.722) |
| nuclear/TME | lymphocyte_density_mm2 | 1522 | 67.7 | 177 | -0.22 | 29 | -0.52 | 2.4e-63 | -0.37 (8.9e-34) |
| nuclear/TME | lymphocyte_fraction | 1522 | 0.0102 | 0.0679 | -0.62 | 18 | -0.81 | 1.3e-155 | -0.76 (8.1e-138) |
| nuclear/TME | neoplastic_fraction | 1522 | 0.852 | 0.0561 | +1.80 | 98 | +1.00 | 3.8e-248 | +1.00 (2.1e-249) |
| nuclear/TME | nuclear_density_mm2 | 1527 | 7.35e+03 | 3.01e+03 | +0.82 | 82 | +0.78 | 1.5e-154 | +0.81 (5.2e-166) |
| artifact | tissue_frac | 1527 | 0.932 | 0.831 | +0.63 | 70 | +0.64 | 1.5e-102 | +0.25 (8.7e-18) |
| artifact | white_frac | 1527 | 0.0682 | 0.169 | -0.63 | 30 | -0.64 | 1.5e-102 | -0.25 (8.7e-18) |
| artifact | blur_laplacian_var | 1527 | 0.00303 | 0.0023 | +0.33 | 79 | +0.70 | 1.7e-123 | +0.72 (1.2e-130) |
| artifact | pen_frac | 1527 | 0.0022 | 0.000622 | +0.05 | 79 | +0.64 | 1.5e-102 | +0.76 (5.9e-142) |
| artifact | dark_frac | 1527 | 0.0169 | 0.00446 | +0.21 | 80 | +0.66 | 1.3e-110 | +0.71 (2.5e-126) |

#### Interpretation notes

- 13/14 primary features differ (q < 0.05). High-attention tiles are tumour-dense with larger, more irregular nuclei and fewer lymphocytes.
- Shaded rows are slide-artifact metrics. If they also differ, attention partly tracks tissue quality; report this and, ideally, repeat the comparison restricted to artifact-free tiles.
- Lymphocytes = HoVer-Net PanNuke 'inflammatory' class; mitoses are not detected. Tiles are ranked within slide from patient-level attention.

### Figure 4. Attention-group differences and attention–IG agreement

Left: forest plot produced by the Aim 2 pipeline (UNI2-h). Right: per-fold agreement between ABMIL attention and |integrated gradients| on held-out patients.

4a. UNI2-h

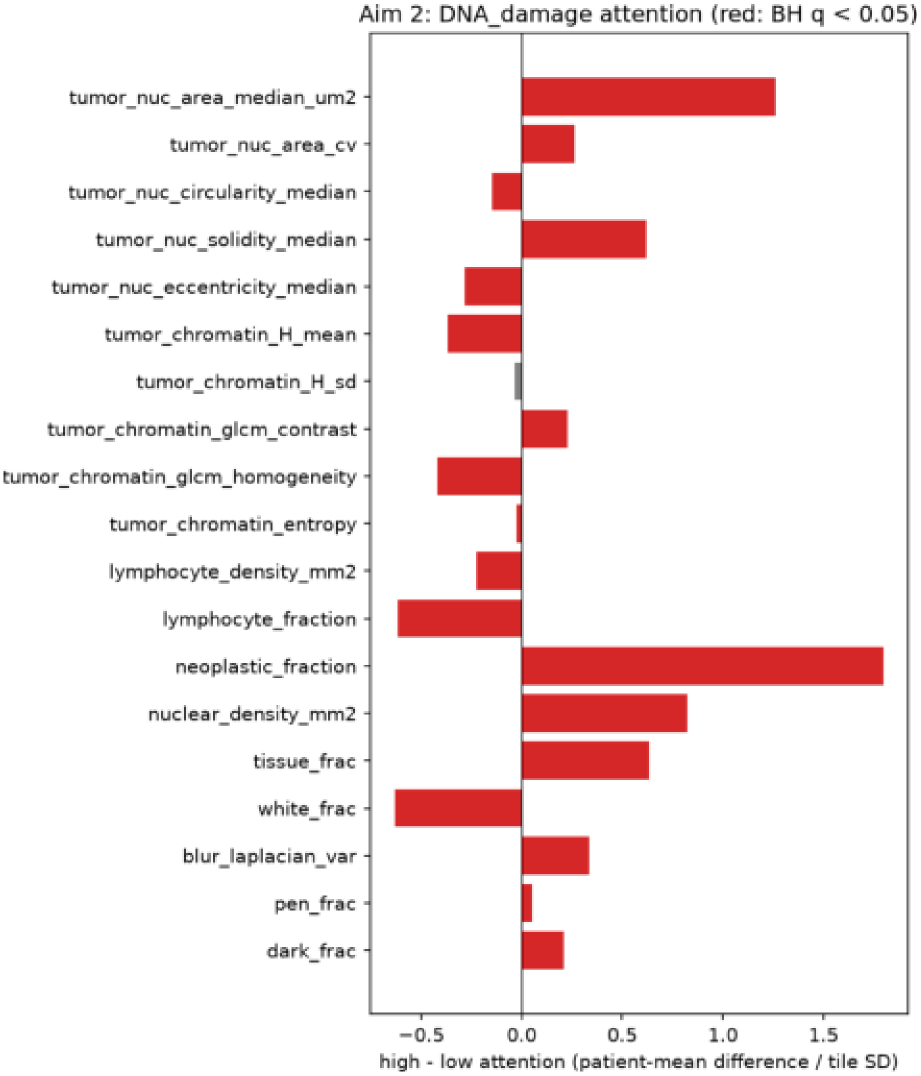


#### 4b. Attention vs IG

| Encoder | Fold | Spearman (att, \|IG\|) | Top-K overlap |
|---|---|---|---|
| UNI2-h | 0 | 0.683 | 0.30 |
| UNI2-h | 1 | 0.731 | 0.30 |
| UNI2-h | 2 | 0.732 | 0.40 |
| UNI2-h | 3 | 0.724 | 0.40 |
| UNI2-h | 4 | 0.662 | 0.30 |
| Virchow2 | 0 | 0.653 | 0.30 |
| Virchow2 | 1 | 0.751 | 0.30 |
| Virchow2 | 2 | 0.705 | 0.20 |
| Virchow2 | 3 | 0.779 | 0.50 |
| Virchow2 | 4 | 0.592 | 0.20 |

#### Attention–IG consistency (UNI2-h, patient level)

- attention vs |IG| on 1527 held-out patients: median Spearman = 0.708 (IQR 0.610-0.776), one-sided Wilcoxon p = 2.00e-251
- top-K overlap median 0.400 vs chance 0.0008, p = 4.12e-250
- signed Spearman(att, IG) median = -0.039
- IG completeness gap: median |gap| = 5.81e-04 (prediction change 3.62e-01)
- Because IG flows through the attention weights, some agreement is built in; present this as a consistency check, not independent validation.

### Figure 5. Pathomic clusters of attended tiles (UNI2-h)

Tiles clustered on HoVer-Net features (k by silhouette). Patient-level median diff: per-patient fraction of high- minus low-attention tiles in the cluster (Wilcoxon, BH q).

| Cluster | Tiles | % high-attention | Defining features (SD from mean) | Patient-level diff median / mean | q (BH) |
|---|---|---|---|---|---|
| 5 | 6,925 | 93 | neopl_fraction +0.8; nuc_area +0.8; nuc_solidity +0.7 | +0.30 / +0.37 | 4.5e-178 |
| 2 | 4,845 | 83 | glcm_contrast +1.6; glcm_homogeneity -1.4; chrom_H_sd +1.0 | +0.00 / +0.18 | 8.8e-99 |
| 1 | 4,877 | 75 | nuc_circularity -1.2; nuc_solidity -1.0; nuc_area_cv +0.9 | +0.00 / +0.14 | 1.1e-76 |
| 3 | 2,770 | 63 | glcm_homogeneity +1.8; chrom_entropy -1.8; chrom_H_mean +1.3 | +0.00 / +0.04 | 3.3e-09 |
| 4 | 1,278 | 9 | lymph_density +3.6; lymph_fraction +2.9; nuclear_density +1.2 | +0.00 / -0.07 | 6.0e-73 |
| 0 | 12,765 | 6 | neopl_fraction -1.0; nuclear_density -0.7; lymph_fraction +0.2 | -0.75 / -0.67 | 8.5e-242 |

Cluster 5 · 93% high-attention

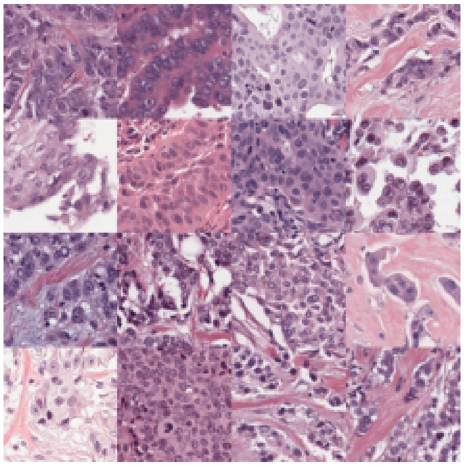

Cluster 2 · 83% high-attention

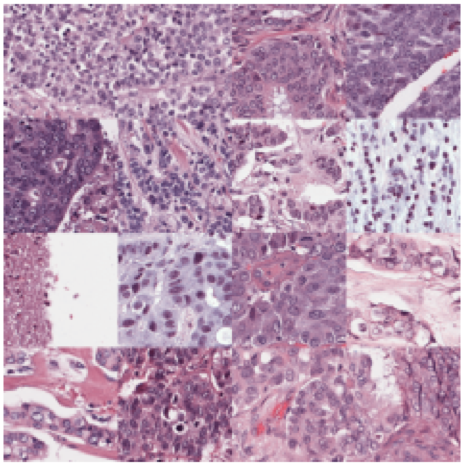

Cluster 1 · 75% high-attention

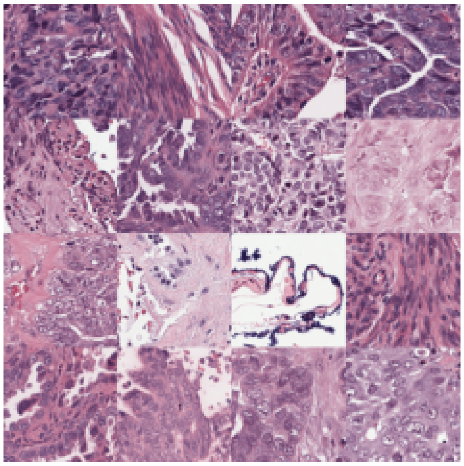

Cluster 3 · 63% high-attention

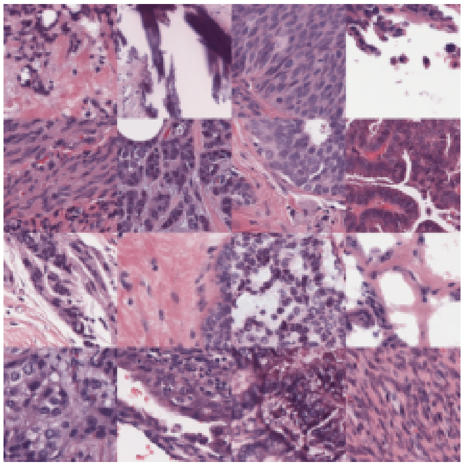

Cluster 4 · 9% high-attention

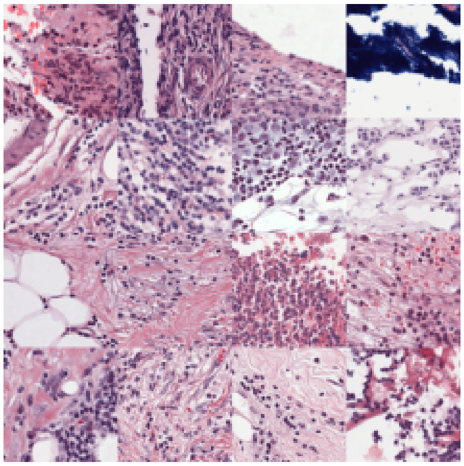

Cluster 0 · 6% high-attention

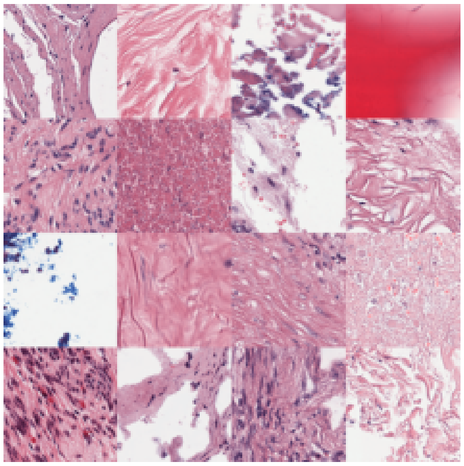

### Figure 6a. Example heatmaps, DNA damage (UNI2-h)

Held-out patients; highest- and lowest-scoring patient per cancer type among slides with heatmaps. Panels: H&E, attention rank (green = top-K, cyan = bottom-K tiles), integrated gradients (red raises the prediction). Values are RPPA pathway scores.

TCGA-AO-A03P · BRCA · measured +0.45, predicted +0.09

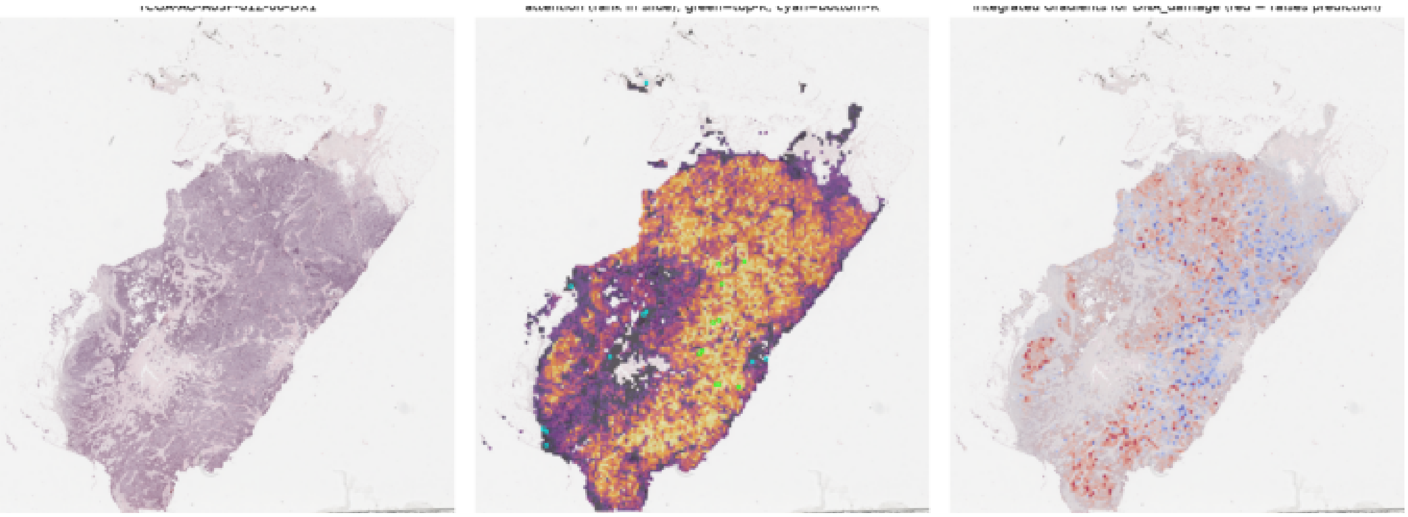

TCGA-E2-A14V · BRCA · measured -0.21, predicted -0.41

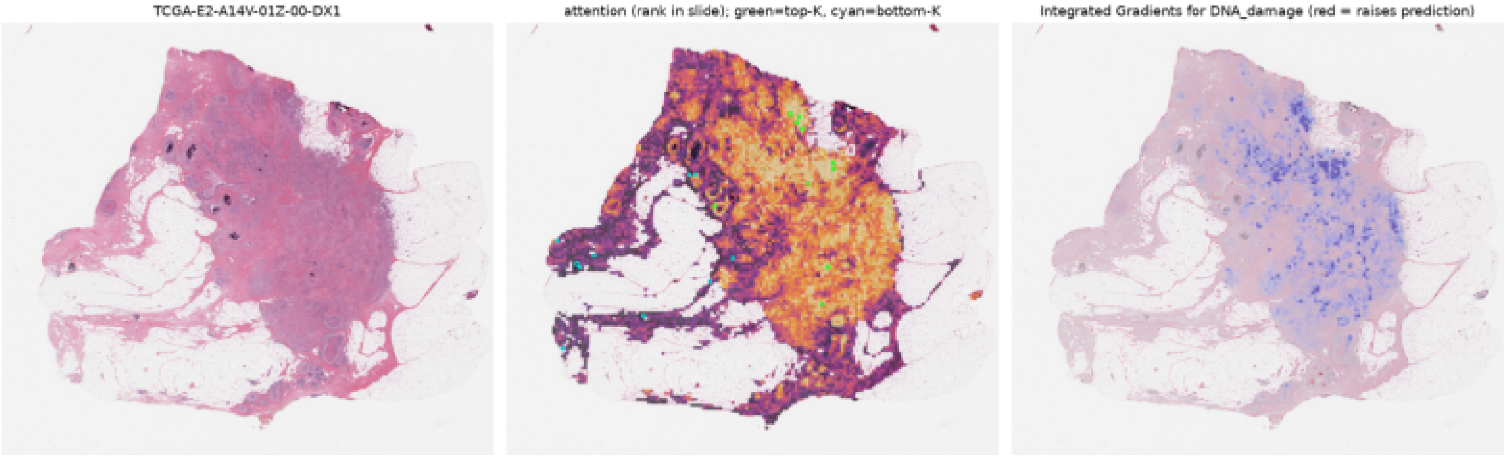


TCGA-BK-A0CB · UCEC · measured +0.05, predicted -0.10

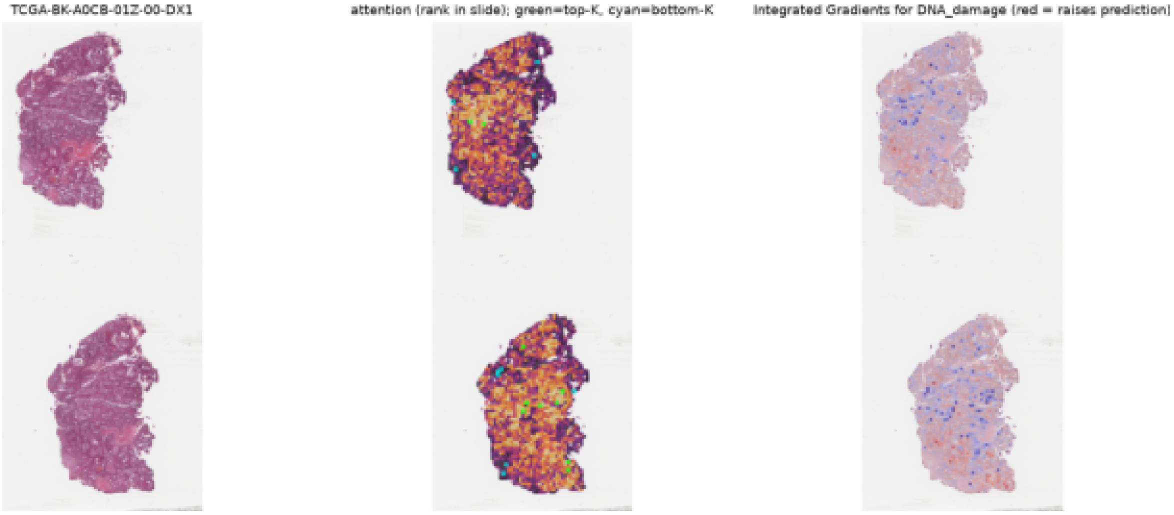

### Figure 6b. Example heatmaps, DNA damage (UNI2-h)

Held-out patients; highest- and lowest-scoring patient per cancer type among slides with heatmaps. Panels: H&E, attention rank (green = top-K, cyan = bottom-K tiles), integrated gradients (red raises the prediction). Values are RPPA pathway scores.

TCGA-AP-A1DM · UCEC · measured +0.05, predicted +0.14

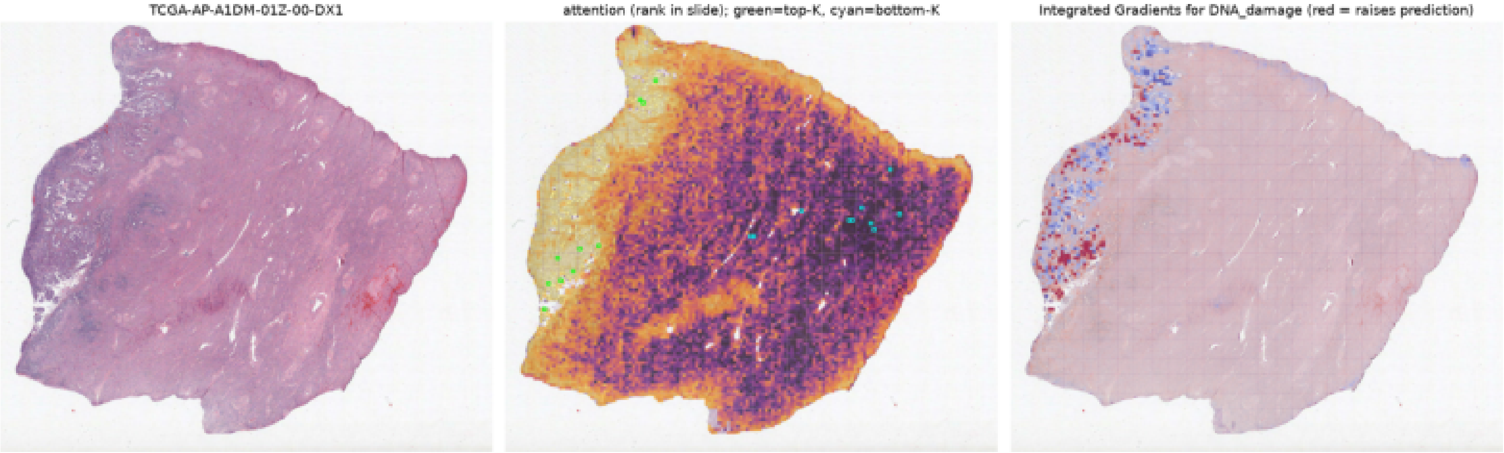


TCGA-4J-AA1J · CESC · measured -0.45, predicted -0.06

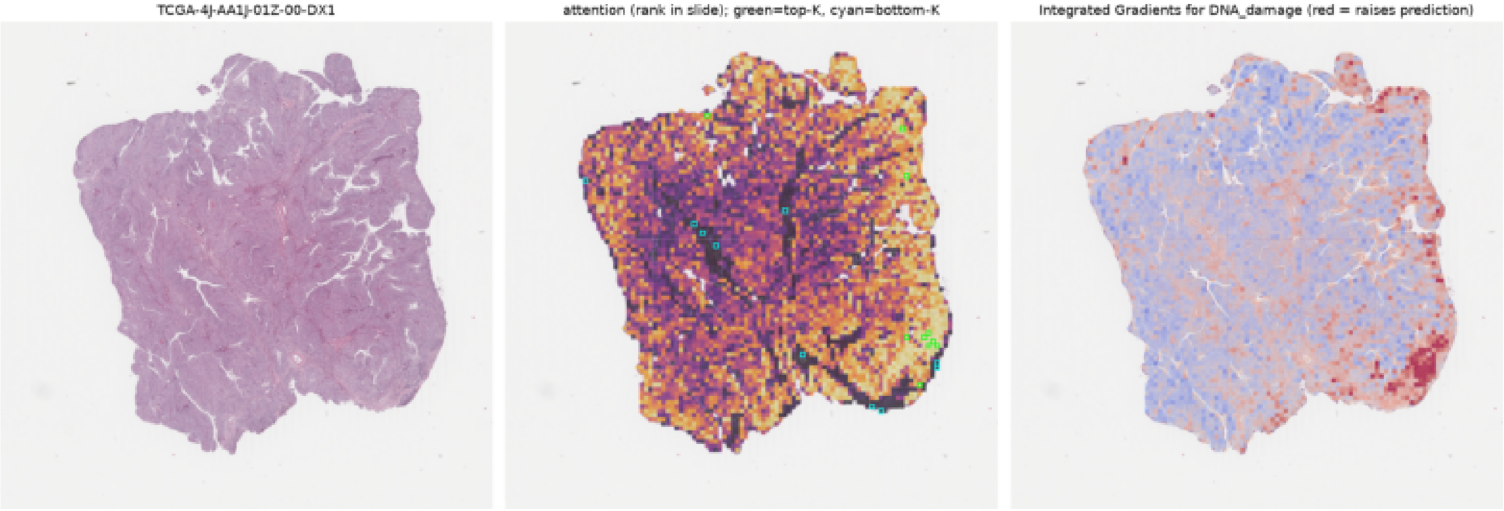

### Figure S2. Aim 2 replication with Virchow2 features

Same tile selection, HoVer-Net features and clustering applied to the Virchow2 model.

S2a. High vs low attention, Virchow2

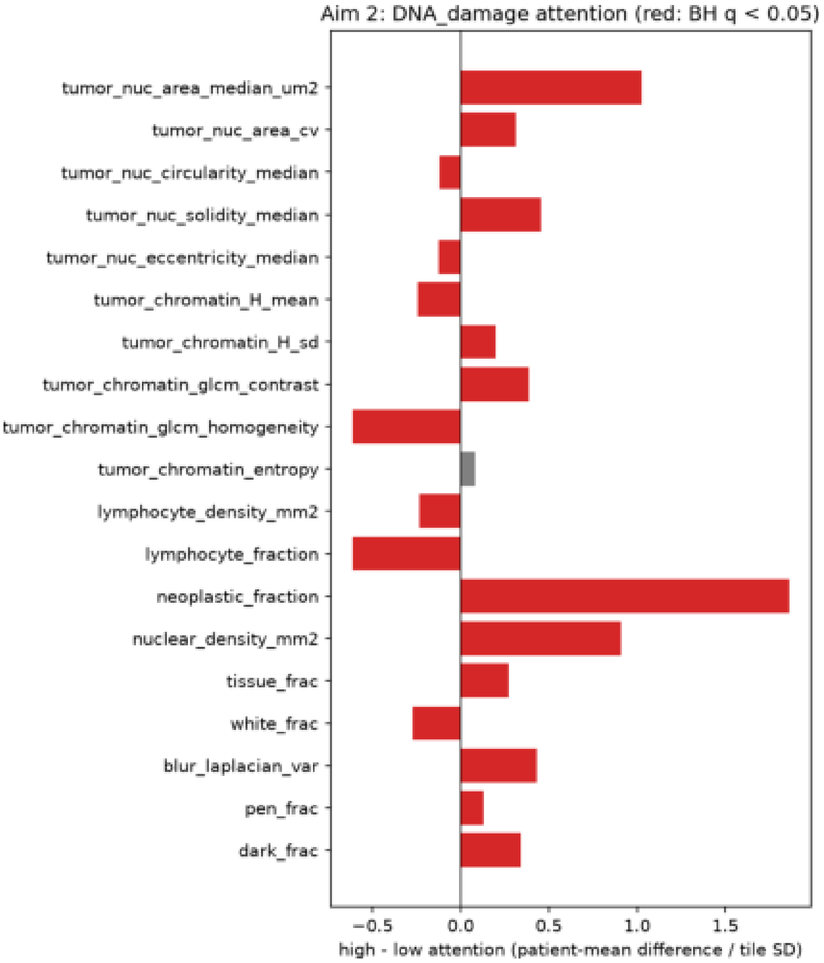


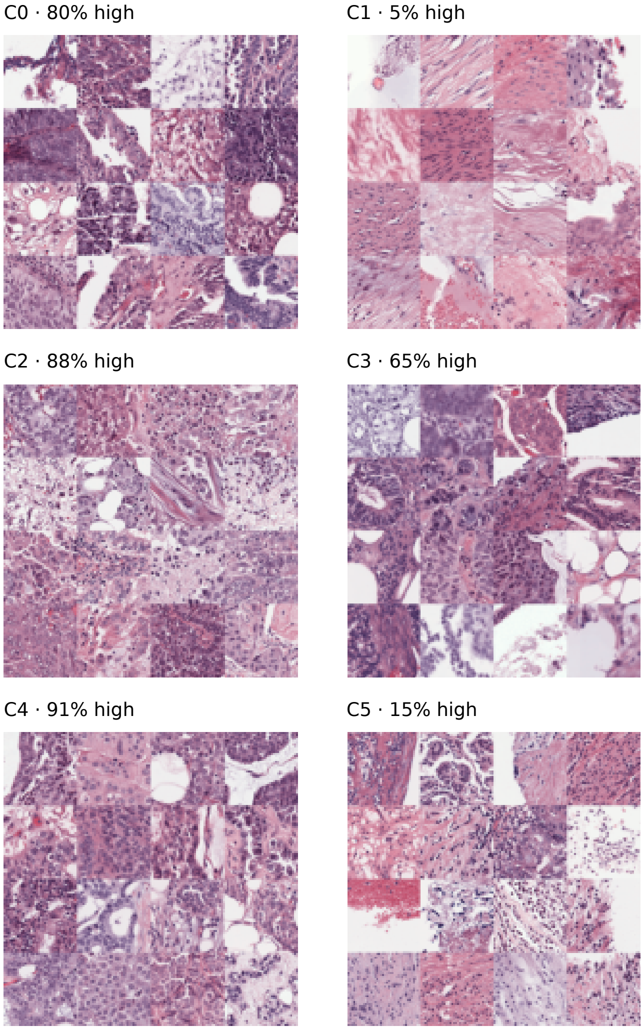


#### Virchow2 Aim 2 summary

- [1] high vs low attention: 13/14 primary features with BH q < 0.05
- [2] attention vs |IG| on 1527 held-out patients: median Spearman = 0.696 (IQR 0.597-0.781), one-sided Wilcoxon p = 2.09e-251
- [3] pathomic clustering: k=6 (silhouette), 6/6 clusters enriched in high vs low attention (patient-level Wilcoxon, BH q < 0.05)
- SUCCESS criterion 1 (nuclear features differ, q<0.05): MET
- SUCCESS criterion 2 (attention agrees with IG on held-out patients; consistency check only: IG flows through the attention weights, so some agreement is built in and this is not independent evidence): MET

### Limitations and items still needed for submission

#### Limitations to state in the manuscript

- Slide and RPPA lysate come from the same tumour sample but not necessarily the same tissue portion; this label noise caps achievable correlation.
- Pathway scores differ strongly by cancer type. Raw $\rho$ partly reflects tissue type, so within-type $\rho$ and $\Delta\rho$ over the type-mean baseline are the claims to lead with.
- p and q values are approximate: pooled CV predictions are not fully independent, and null targets drift slightly negative under CV. Report bootstrap CIs and fold-level sign consistency alongside them.
- OV ($n \approx 72$ with DX slides) and UCS ($n \approx 48$) are small; per-type estimates there are imprecise.
- RPPA has no direct ATR, DNA-PKcs, WEE1 or γH2AX readouts; DNA damage response activity is a composite protein score, not a direct kinase-activity assay. PARP1 is largely missing and was not used.
- Aim 2 artifact metrics also differ between attention groups, so attention partly tracks tissue quality.
- All results are internal TCGA cross-validation.

#### Not in this results set (typically requested by reviewers)

- External validation on an independent cohort (e.g., CPTAC BRCA/UCEC/OV with matched proteomics), or an explicit statement that it is out of scope.
- Simple comparison baselines on the same folds: mean-pooled features + ridge regression, and/or CLAM or TransMIL, to show what ABMIL adds.
- A statement on multiple-seed variability (all runs use seed 0).
- Final-figure versions at journal resolution (vector PDF or ≥300 dpi TIFF) once the figure set is fixed.
- Data and code availability statement (GDC/TCGA accessions, RPPA source file, repository link).

### Reproducibility record

Source directory: /projects/wgs_pipeline_dev/abmil_runs. Tables on these pages are also written as CSV to /projects/wgs_pipeline_dev/paper_report/tables.

| Label | Run dir | Features | Bags file | Folds | Target mode | Shuffled | Best epochs |
|---|---|---|---|---|---|---|---|
| UNI2-h | uni2h_11-3-26 | 1536-d | abmil_data_uni2h/bags_unifolds.csv | file | global | no | {0:6,1:5,2:5,3:7,4:3} |
| Virchow2 | virchow2_11-3-26 | 2560-d | abmil_data_virchow2/bags_unifolds.csv | file | global | no | {0:7,1:8,2:4,3:7,4:3} |
| UNI (v1) | 11-2-26 | 1024-d | abmil_data_uni/bags.csv | default | global | no | {0:5,1:7,2:5,3:5,4:4} |
| UNI, site-stratified CV | uni_debias | 1024-d | abmil_data_uni/bags.csv | site | residual | no | {0:6,1:5,2:5,3:7,4:3} |
| UNI2-h null | uni2h_pathways_null | 1536-d | abmil_data_uni2h/bags_unifolds.csv | file | global | yes | {0:3,1:3,2:3,3:2,4:3} |
| Virchow2 null | virchow2_pathways_nul | 2560-d | abmil_data_virchow2/bags_unifolds.csv | file | global | yes | {0:5,1:3,2:1,3:1,4:3} |
| UNI site-strat. null | uni_debias_null | 1024-d | abmil_data_uni/bags.csv | site | residual | yes | {0:2,1:2,2:1,3:1,4:1} |

#### Pipeline scripts

- 01_download_gdc.sh, 02_match.py, 03_normalize.py, 03_normalize.sbatch, 04_qc_sample.py, 05_train.py, 05_train.sbatch, 06_metrics.py, 07_aim2_explain.py, 07_aim2_submit.sh, 07_fix_coords.py, fm_extract.sbatch, fm_train.sbatch, fm_train_null.sbatch, run_pipeline.sbatch, train_debias.sbatch, train_uni.sbatch, uni_extract.sbatch

#### Other runs not reported here

- 10-23-26 and 10-28-26 are early pilot runs on the original feature set (abmil_data/bags.csv) and are superseded by the runs above.

#### Software

- Report built with Python 3.11.17, pandas 3.0.6, matplotlib 3.11.2, numpy 2.4.6.
